# A role for gene expression and mRNA stability in nutritional compensation of the circadian clock

**DOI:** 10.1101/2022.05.09.491261

**Authors:** Christina M. Kelliher, Elizabeth-Lauren Stevenson, Jennifer J. Loros, Jay C. Dunlap

## Abstract

Compensation is a defining principle of a true circadian clock, where its approximately 24-hour period length is relatively unchanged across environmental conditions. Known compensation effectors directly regulate core clock factors to buffer the oscillator’s period length from variables in the environment. Temperature Compensation mechanisms have been experimentally addressed across circadian model systems, but much less is known about the related process of Nutritional Compensation, where circadian period length is maintained across physiologically relevant nutrient levels. Using the filamentous fungus *Neurospora crassa*, we performed a genetic screen under glucose and amino acid starvation conditions to identify new regulators of Nutritional Compensation. Our screen uncovered 16 novel mutants, and together with 4 mutants characterized in prior work, a model emerges where Nutritional Compensation of the fungal clock is achieved at the levels of transcription, chromatin regulation, and mRNA stability. However, eukaryotic circadian Nutritional Compensation is completely unstudied outside of *Neurospora*. To test for conservation in cultured mammalian cells, we selected the top two hits from our fungal genetic screen, performed siRNA knockdown experiments of the mammalian homologs, and characterized the cell lines with respect to compensation. We find that the wild-type mammalian clock is also compensated across a large range of external glucose concentrations, as observed in *Neurospora*, and that knocking down *CPSF6* or *SETD2* in human cells also results in nutrient-dependent period length changes. We conclude that, like Temperature Compensation, Nutritional Compensation is a conserved circadian process in fungal and mammalian clocks and that it may share common molecular determinants.

## Introduction

Circadian clocks exist at the cellular level to allow cell types, tissues, and organisms to properly align physiology with time of day. True circadian clocks are sensitive to the external environment in two distinct ways. Discrete pulses of bright light, temperature, nutrients, hormones, or other chemicals reset circadian oscillators and re-orient the clock’s phase to the new environment (reviewed in: Johnson et al., 2003). This resetting feature of circadian clocks is most commonly experienced when jetlagged humans travel across multiple time zones and entrain to the destination’s light/dark cycles. On the other hand, circadian clocks are also shielded or buffered from changes in the ambient environment within the physiological range of an organism (Hastings and Sweeney, 1957; Pittendrigh et al., 1959). In the filamentous fungus *Neurospora crassa*, circadian period length is maintained at approximately 21.5 hours when grown at constant temperatures ranging from 16°C to 32°C (Gardner and Feldman, 1981) or under different nutrient conditions (Sargent and Kaltenborn, 1972). This circadian property is known as compensation, and the molecular mechanisms underlying period length compensation remain elusive.

At the molecular level, the circadian oscillator is a transcription-translation feedback loop (TTFL) that is functionally conserved from fungi to animals. The positive arm of the clock is composed of transcription factor activators (WC-1/WC-2 in fungi; BMAL1/CLOCK in mammals), which form a heterodimeric complex, drive transcription of direct target genes, and recruit chromatin modifiers (Crosthwaite et al., 1997; Koike et al., 2012; Menet et al., 2014). The positive arm directly regulates the negative arm of the clock (FRQ in fungi; PERs/CRYs in mammals) (Aronson et al., 1994; reviewed in: Dunlap, 1999; Philpott et al., 2021). Negative arm clock components form a stable complex with Casein Kinase I (CKI) and other factors, leading to feedback and posttranslational inhibition of the positive arm to close the circadian feedback loop (Wang et al., 2019; Cao et al., 2021). The positive and negative arms are sufficient for rhythmicity, although accessory feedback loops confer additional clock robustness (reviewed in: Takahashi, 2017). Such individual cellular clocks are coordinated in a coupled network to align organismal physiology in mammals (reviewed in: Finger et al., 2020).

Temperature Compensation was first proposed to be a circadian property in the dinoflagellate *Gonyaulax polyedra* when increasing temperatures led first to period lengthening (so-called “over-compensation”) and then to period shortening at even higher temperatures, a result that plainly conflicted with models based on biochemical reaction rates strictly increasing with temperature (Hastings and Sweeney, 1957). In genetic model systems like *Neurospora* and *Drosophila* where organisms operate at ambient temperatures, and even in homeothermic animals, cellular circadian clocks are temperature compensated (Zimmerman et al., 1968; Gardner and Feldman, 1981; Barrett and Takahashi, 1995; Izumo et al., 2003; Tsuchiya et al., 2003; Kidd et al., 2015). The forward and reverse genetics that have driven current models for Temperature Compensation have led to casein kinases as central regulators across multiple circadian model systems. In *Neurospora* and in plants, Casein Kinase II (CKII) is required for Temperature Compensation (Mehra et al., 2009; Portolés and Más, 2010). *Neurospora* CKII activity increases linearly with temperature and directly phosphorylates the negative arm of the clock (Mehra et al., 2009). The *tau* mutant hamster (CKIε^R178C^) was the first characterized mammalian Temperature Compensation defect (Ralph and Menaker, 1988; Tosini and Menaker, 1998; Lowrey et al., 2000). Recent structural work has demonstrated that CKI *tau* alters both priming and progressive phosphorylation events on the negative arm of the clock (Philpott et al., 2020), which may account for its period shortening with temperature (so-called “under-compensation”). Indeed, temperature-sensitive target binding has been shown for mammalian CKI (Shinohara et al., 2017), and the interaction strength between CKI and FRQ in *Neurospora* has been implicated in Temperature Compensation as well (Hu et al., 2021). Taken together, CKI and CKII each have temperature sensitive aspects of enzyme activity, both can directly phosphorylate core clock components, and casein kinases appear to play a conserved role in Temperature Compensation of fungal, plant, and animal clocks.

In contrast to Temperature Compensation, mechanisms underlying Nutritional Compensation (also known as Metabolic or Glucose Compensation) are poorly understood in *Neurospora* and completely unstudied in other eukaryotic circadian systems. Period compensation to a variety of ATP:ADP ratios has been well described in the prokaryotic cyanobacteria *Synechococcus elongatus* (reviewed in: Johnson and Egli, 2014). A handful of Nutritional Compensation defects have arisen sporadically in *Neurospora*, the most developed involving the transcription factor repressor CSP-1 (Lambreghts et al., 2009). CSP-1 directly regulates and is regulated by the clock’s positive arm White Collar Complex (WCC), forming an accessory negative feedback loop. In a Δ*csp-1* mutant, period significantly shortens as a function of glucose concentration (Sancar et al., 2012). In fact, direct overexpression of *wc-1* also causes nutritional under-compensation (Dovzhenok et al., 2015). Nutritional Compensation is defective in the absence of the general transcription repressor RCO-1 (Olivares-Yanez et al., 2016), likely due its normal role in preventing WCC-independent *frq* transcription (Zhou et al., 2013). Over-compensation was also found in loss-of-function mutants of an RNA helicase, PRD-1, which normally localizes to the nucleus only under high glucose conditions (Emerson et al., 2015). Nutrient sensing and signaling pathways should presumably also play a role in Nutritional Compensation of the clock, and RAS2 and cAMP signaling have been implicated (Gyöngyösi et al., 2017). Taken together, the current incomplete model for Nutritional Compensation in *Neurospora* assembled from random hits implicates transcriptional and post-transcriptional regulation of core clock factors by transcription factors, an RNA helicase, and cAMP signaling.

Since the realization of circadian compensation, the field has speculated that Temperature and Nutritional Compensation pathways may share common regulators (Pittendrigh and Caldarola, 1973; Roenneberg and Merrow, 1999; Johnson and Egli, 2014). We directly test this model and find in *Neurospora* that previously reported compensation mutants are specific to either Temperature or Nutritional Compensation. Given this separation of function and relatively little mechanistic knowledge about Nutritional Compensation, we designed a genetic screen to identify new compensation mutants in *Neurospora crassa*. We identify 16 gene knockouts with Nutritional Compensation phenotypes, greatly expanding the list of 4 previously characterized mutants. We also provide the first evidence that, like Temperature Compensation, Nutritional Compensation is relevant in the mammalian circadian system, and confirm that knockdowns of the human genes homologous to the top two hits from our genetic screen, *CPSF6* and *SETD2*, impair the clock’s ability to maintain its period length at different glucose levels. These data establish a potentially conserved genetic basis for the phenomenon of circadian Nutritional Compensation and anchor the phenomenon for further genetic and molecular dissection.

## Results

### Nutritional Compensation is distinct from Temperature Compensation in *Neurospora*

We first set out to characterize the properties of Nutritional Compensation in wild-type *Neurospora*. Traditional circadian experiments have utilized the *ras-1^bd^* mutant, which promotes the formation of circadianly-regulated distinct bands of conidial spores in a race tube assay (Sargent et al., 1966; Belden et al., 2007a). However, the *ras-1* gene (NCU08823) is implicated in growth and regulation of reactive oxygen species. Thus to accurately profile normal Nutritional Compensation, a wild-type *ras-1^+^* strain containing a *frq* clock box transcriptional reporter was used to measure period length across glucose concentrations ranging from 0 –111 mM (0 – 2% w/v). The *Neurospora* circadian period is slightly under-compensated to nutrients (Figure 1A); under-compensation has also been observed with respect to temperature (Mehra et al., 2009; Hu et al., 2021). Fungal biomass increases by orders of magnitude when grown in the range of 0% – 0.75% w/v glucose (Supplementary Figure 1), and this increased biomass accounts for the increased magnitude of luciferase rhythms observed at higher glucose levels (Figure 1B) (Supplementary Movie 1).

**Figure 1.**
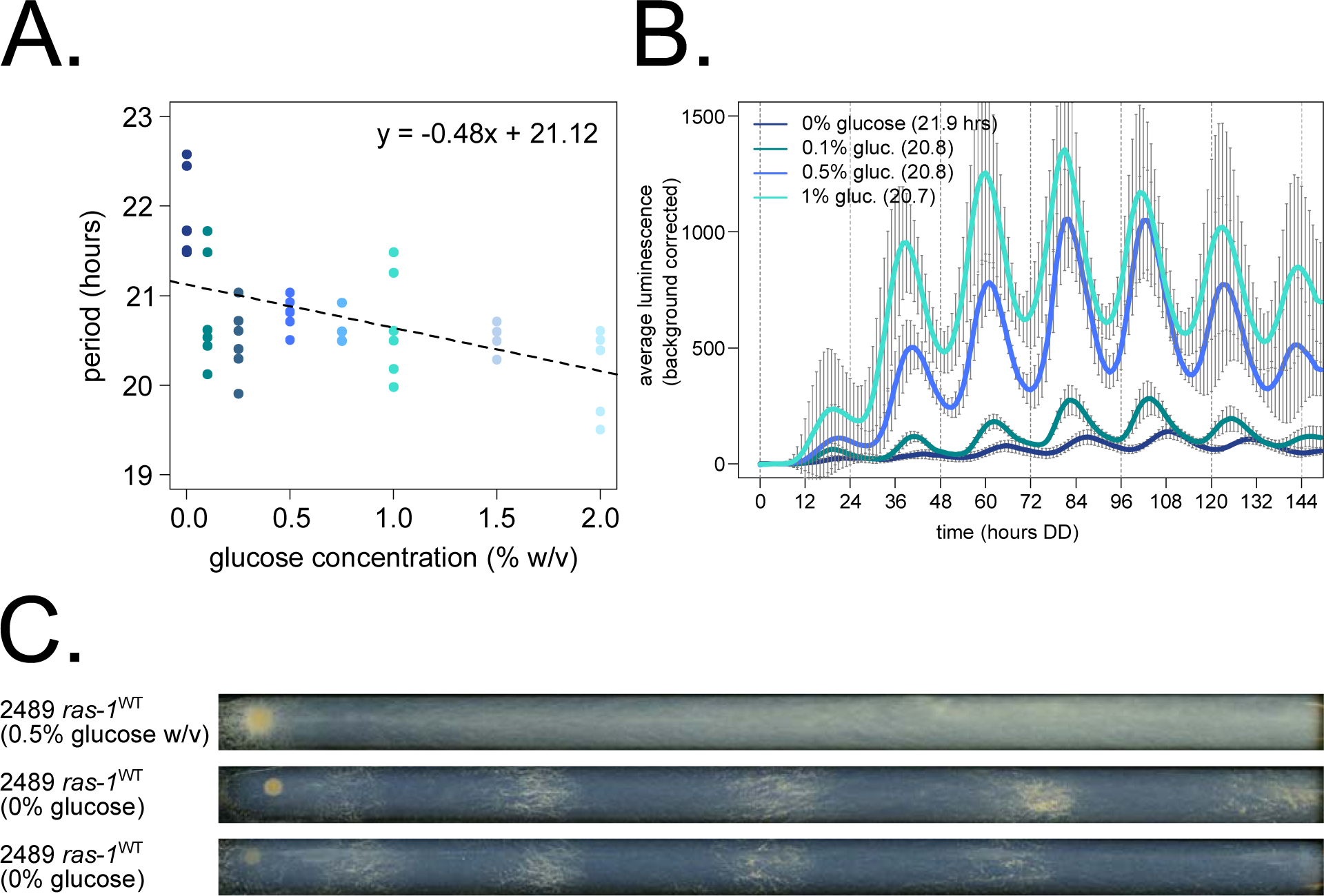
Nutritional Compensation properties of *Neurospora crassa*. Circadian period length was determined by bioluminescent recordings of a *ras-1^WT^* strain cultured on race tubes, where the fungal growth front encounters constant glucose concentrations (N = 6 race tubes per concentration). No arginine was added to the medium. Period shortens slightly as a function of glucose levels, which is indicated by the negative slope of the linear fit (`glm` in R, Gaussian family defaults, slope = –0.48 ± 0.12) (**A**). Averaged biological and technical replicates are shown for 0 mM (0% w/v), 5.6 mM (0.1% w/v), 27.8 mM (0.5% w/v), and 55.5 mM (1% w/v) glucose levels (standard deviation error bars). Period lengths are 21.9 ± 0.5, 20.8 ± 0.6, 20.8 ± 0.2, and 20.7 ± 0.6 hours, respectively (average ± SD) (**B**). Surprisingly, the *ras-1^WT^* strain can form distinct conidial bands when grown on 0% glucose starvation medium, contrasted with constitutive conidiation seen at high glucose levels (**C**).

Having established that the *Neurospora* clock displays compensation for period length across glucose concentrations, we asked whether Temperature Compensation mutants also have Nutritional Compensation phenotypes, and vice versa, together in the same assay. CKII is required for normal Temperature Compensation, and its catalytic subunit mutant *cka^prd-3^* (Y43H) has an over-compensation phenotype by race tube assay (Mehra et al., 2009). The *frq^7^* (G459D) point mutant is temperature under-compensated (Gardner and Feldman, 1981). The *frq* clock box transcriptional reporter was integrated into the *cka^prd-3^* and *frq^7^* mutant backgrounds, and temperature over- (Q_10_ = 0.88) and under-compensation (Q_10_ = 1.11) phenotypes were confirmed by luciferase assay (Figure 2A). Known Nutritional Compensation mutants *prd-1* and Δ*csp-1* were also tested, and each had normal Temperature Compensation profiles (Q_10_ = 1.04, 1.06). The same 4 compensation mutant reporter strains were then grown on race tubes containing zero or high glucose medium, and bioluminescence was recorded to track period length and Nutritional Compensation phenotypes. Controls were slightly under- compensated (Figure 1A, Figure 2B). Nutritional Compensation mutants *prd-1* and Δ*csp-1* showed the over- and under-compensation phenotypes reported by previous studies (Sancar et al., 2012; Emerson et al., 2015). Temperature Compensation mutants *cka^prd-3^* and *frq^7^* have normal Nutritional Compensation (Figure 2B). We conclude that Temperature and Nutritional Compensation are controlled by distinct pathways in *Neurospora*. These data suggest that further examination of available mutants defective in Temperature Compensation will not inform our understanding of Nutritional Compensation, and that a separate genetic screen is warranted.

**Figure 2.**
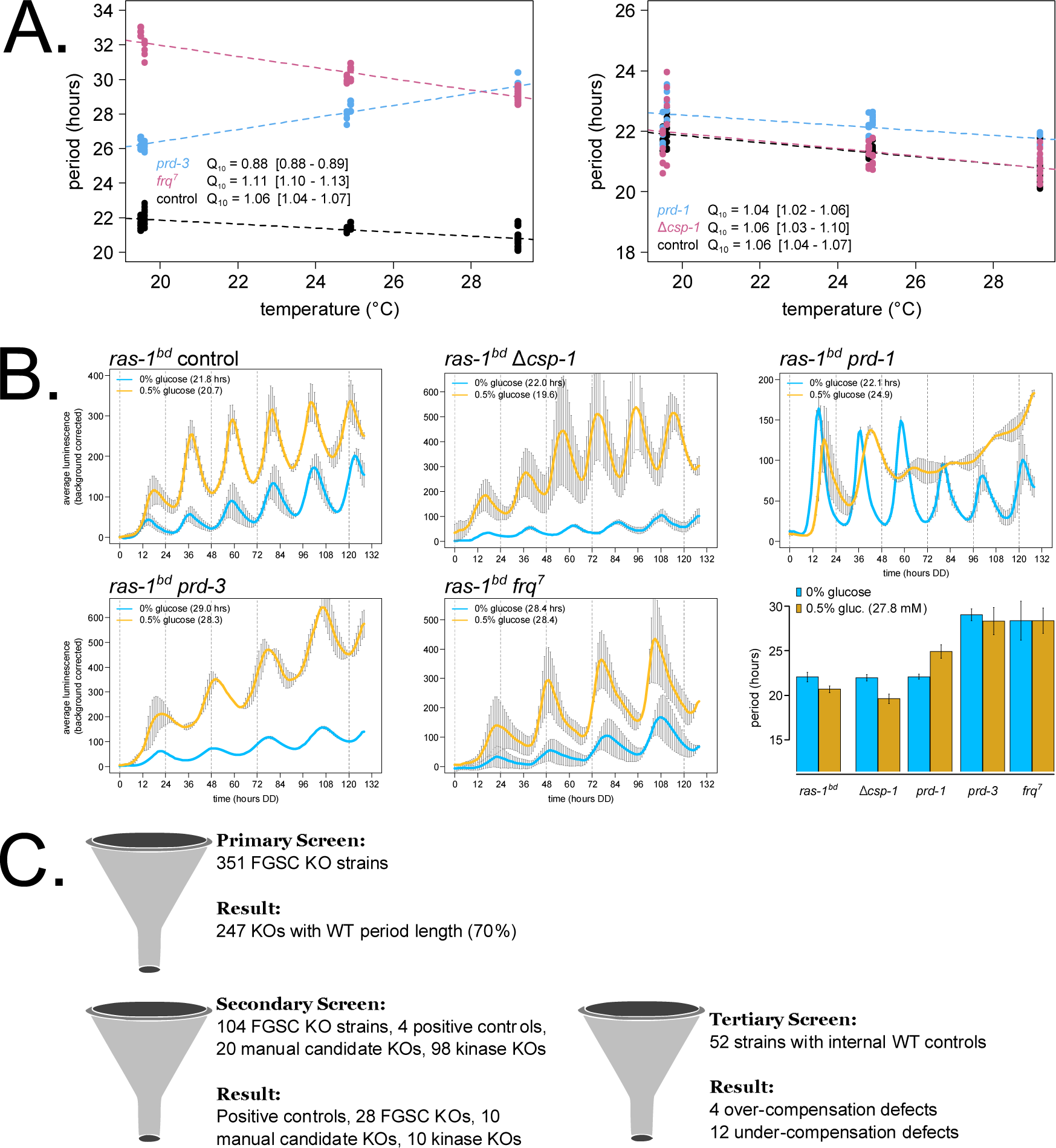
Nutritional Compensation mutants have normal Temperature Compensation in *Neurospora*, catalyzing a genetic screen for new mutants. 96-well plate luciferase assays were used to measure the circadian period length across temperatures in constant darkness (N = 12 replicates per strain, per temperature). A linear model was fit to period length data from each strain (`glm` in R), and a Q_10_ temperature coefficient was calculated using the model-fitted period lengths at 20°C and 30°C. Shaded areas around the linear fit represent the 95% confidence intervals on the slope. Error ranges on Q_10_ values were computed from the 95% CIs (**A**). Circadian bioluminescence was recorded from race tube cultures of the indicated genotypes, as previously described (Larrondo et al., 2012) (Supplementary Movie 1). High nutrient medium (yellow lines) contained 0.5% w/v glucose 0.17% w/v arginine, and zero nutrient medium (blue lines) contained 0% glucose 0% arginine. Period lengths were computed (N = 2 – 5 biological replicates per nutrient concentration) and summarized in a bar graph (**B**). Cartoon depiction of the 3-phase genetic screen for Nutritional Compensation defects among kinase, RNA regulatory proteins, and manual candidate knockout strains (**C**).

To achieve this, we leveraged the whole genome knockout collection in *Neurospora crassa* (Colot et al., 2006) to initiate a screen for new Nutritional Compensation regulators. Two major classes of gene knockouts were selected to search for compensation phenotypes. Kinases are central to many aspects of cellular processes and regulation including responses to the environment (pheromones, osmotic conditions, carbon/nitrogen regulation, etc.), and a collection of ∼100 different kinase knockout circadian reporter strains was available from previous work (Dasgupta, 2015). In addition to posttranslational modifications by kinases, posttranscriptional regulation is emerging as critically important for circadian output (Hurley et al., 2018). Together with the dramatic nutritional over-compensation defect seen in *prd-1* RNA helicase mutants (Emerson et al., 2015) (Figure 2B), we selected 351 putative RNA regulatory protein knockouts to screen for compensation regulators. Our list of putative RNA-binding and RNA regulatory proteins was derived from bioinformatic databases and from the literature (Ray et al., 2013; Hogan et al., 2015; Zaveri et al., 2017; Basenko et al., 2018) to include proteins with nucleotide-binding functional annotation but exclude known transcriptional regulators. Multiple circadian period alterations were identified in a recent characterization of transcription factor knockouts (Muñoz-Guzmán et al., 2021), and Nutritional Compensation defects among transcription factors, in addition to CSP-1 and RCO-1, will be the subject of future study. Finally, a handful of manually selected candidate genes were included in the compensation screen. 8 classical alleles with long or short circadian period (the series of *prd* mutants and *frq* point mutants) were included, and 12 knockouts were manually selected based on reported roles in nutrient sensing and signaling.

The genetic screen for Nutritional Compensation defects was divided into three phases (Figure 2C). The 351 previously uncharacterized RNA regulatory protein knockout strains were grown on glucose starvation race tubes (Figure 1C) to identify those with period length differences ±1 hour as compared to internal wild-type controls (N = 2 – 8 replicate race tubes per KO strain). Approximately 70% of the putative RNA regulatory protein knockout strains showed normal period length on starvation medium and were eliminated in the primary screen (Figure 2C, Supplementary Table 1). During this phase of the screen, 7 knockout strains were found with enhanced conidial banding patterns (and variable growth rates) relative to wild-type controls, reminiscent of the *ras-1^bd^* mutant phenotype (Supplementary Figure 2). In the secondary screen, the *frq* clock box transcriptional reporter was integrated into all knockouts of interest, along with the existing kinase deletion collection (Dasgupta, 2015) and manually selected candidate strains. This collection of more than 200 strains was screened in a high throughput 96-well plate format using glucose and amino acid starvation medium to identify period length differences ±1 hour from internal wild-type controls (N = 6 – 12 replicate wells per KO strain). About 77% of the candidate strains were eliminated during the secondary screen due to normal period length on starvation medium (Figure 2C, Supplementary Table 1). The ∼50 remaining knockout strains and wild-type controls were then advanced to the lowest throughput tertiary screen: bioluminescence race tube assays directly comparing zero versus high glucose and amino acid medium (0.5% w/v glucose, 0.17% w/v arginine). It should be noted here that the race tube assay is particularly well suited for a Nutritional Compensation mutant screen because the growth front, which produces most of the bioluminescence, always encounters fresh medium; thus there is no complicating effect of nutrient depletion over the course of a six-day assay (Supplementary Movie 1). Circadian period lengths were quantified to assay Nutritional Compensation phenotypes. 16 new mutants emerged from the genetic screen showing large period changes between zero and high nutrient conditions (Figure 2C, Supplementary Table 1).

### mRNA stability and polyadenylation factors emerge as key regulators of circadian period length across nutrients

We first examined the group of 12 under-compensation mutants, where period length shortens as a function of nutrient levels (Figure 3, Supplementary Table 1). The two most significant hits are subunits of the nonsense-mediated decay machinery in *Neurospora* and showed more dramatic nutrient-responsive period shortening than Δ*csp-1* (Sancar et al., 2012). Additionally, 3 of the 12 mutants share a common function in polyadenylation of nascent RNAs. NCU02736 (FGSC12857) is designated as an uncharacterized gene in *Neurospora* but is broadly conserved in fungi. Its *Saccharomyces cerevisiae* homolog is a component of the mRNA cleavage and polyadenylation factor I complex (YGL044C, *RNA15*). PABP-2 (NCU03946, FGSC19900) binds in poly(A) tail regions and can broadly regulate mRNA stability. NAB2 (NCU16397, FGSC22799^Het^) shares a common domain with the yeast gene *NAB2* (YGL122C), which plays a role in mRNA export and stability. Taken together, regulation of polyadenylation and/or mRNA stability is a common axis of period maintenance with increasing nutrients. The other 6 under-compensation mutants did not share an obvious functional pathway, although some of these genes do show circadian rhythms at the mRNA or protein level and/or are regulated in response to light (Supplementary Table 2). However, given the slight under-compensation phenotype described for the wild-type clock in *Neurospora* (Figure 1A, Figure 2B, Figure 3), the remaining under-compensation mutants were not pursued further.

**Figure 3.**
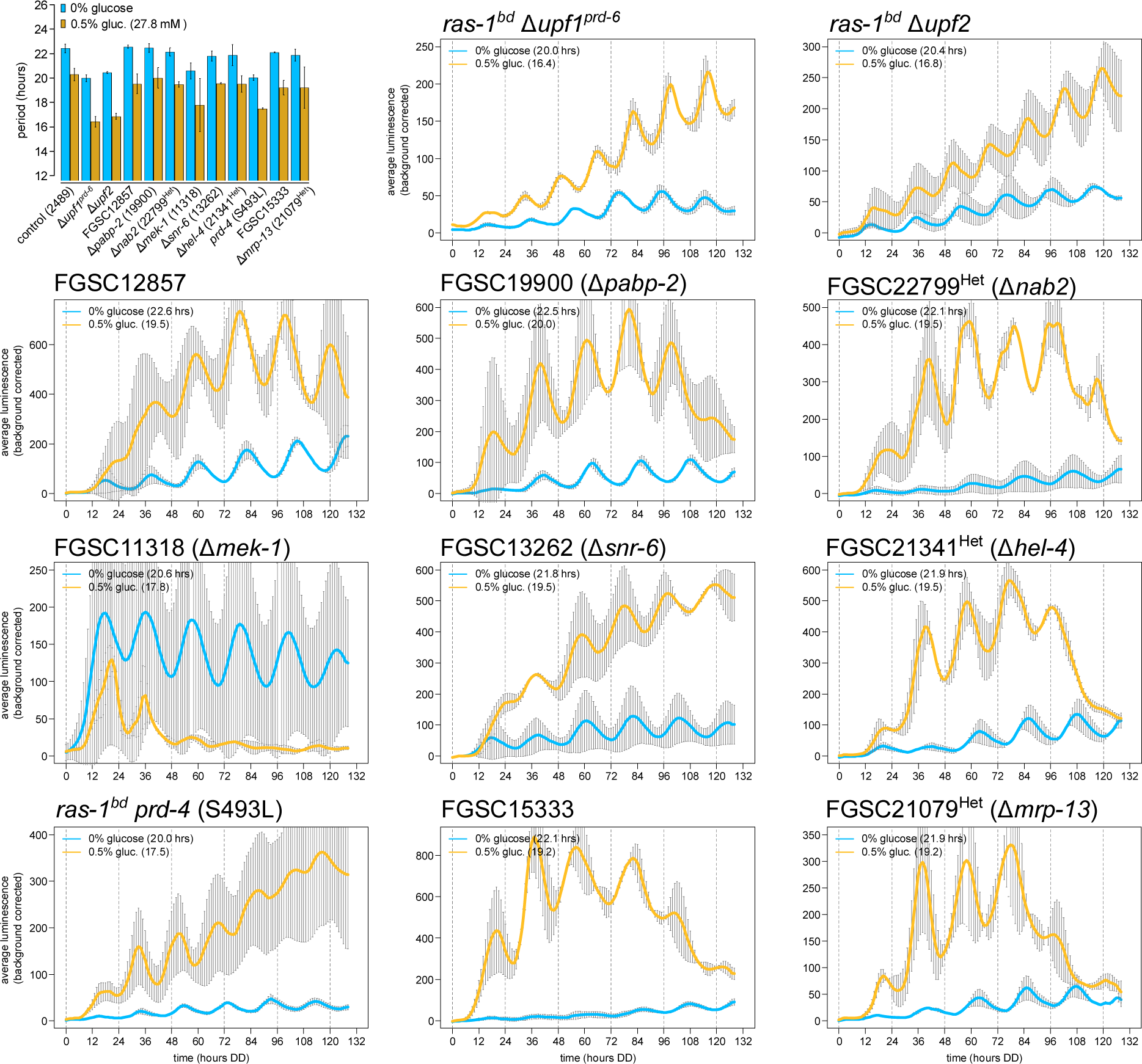
Nutritional under-compensation mutants are enriched for regulators of mRNA stability. Circadian bioluminescence was recorded from race tube cultures of the indicated deletion mutants. High nutrient medium (yellow lines) contained 0.5% w/v glucose 0.17% w/v arginine, and zero nutrient medium (blue lines) contained 0% glucose 0% arginine. Period lengths were computed (N ≥ 2 biological replicate period estimates per nutrient concentration) and summarized in a bar graph compared to controls. “Het” indicates heterokaryon strains derived from the *Neurospora* whole genome deletion collection, which were maintained on hygromycin selection medium prior to the bioluminescence race tube assays.

The nonsense-mediated decay (NMD) pathway is required for a normal circadian period length due to its regulation of *casein kinase I* mRNA levels (Kelliher et al., 2020a). We hypothesized that NMD regulation of *ck-1a* also underlies its nutritional under-compensation defect. NMD in *Neurospora* can be triggered by long 3’ UTR sequences, and the *ck-1a* 3’ UTR is among the longest 1% of UTRs in *Neurospora* (Kelliher et al., 2020a). We generated a *ck-1a* mutant strain lacking its entire 3’ UTR, decoupling *ck-1a* transcripts from NMD targeting. The *ck-1a*Δ3’UTR strain does indeed have a short period length phenocopying that of NMD mutants; however, the *ck-1a*Δ3’UTR under-compensation phenotype is not as severe as observed in NMD mutants (Supplementary Figure 3). This suggests that multiple NMD targets are required for normal Nutritional Compensation. Interestingly, NMD mutants also show ∼1.4-fold changes in *wc-2* and *frh* gene expression levels (Kelliher et al., 2020a: Figure 5B; Wu et al., 2017). *wc-2* is upregulated in NMD mutants, which could easily explain the under-compensation phenotype because *wc-1* overexpression alone is sufficient to drive nutritional under-compensation (Dovzhenok et al., 2015). Curiously, we find that *frh*, the obligate binding partner of the disordered protein FRQ (Hurley et al., 2013), is 18-fold down-regulated in response to carbon starvation in wild-type *Neurospora* (Wang et al., 2017) (Supplementary Figure 4). In fact, both *prd-1* and *frh* are among the top 220 genes in the entire *Neurospora* transcriptome that decrease in expression after glucose starvation. Future work will determine whether NMD regulation of *ck-1a*, *wc-2*, *frh*, or all transcripts explains the nutritional under-compensation defect.

Mutations in two genes, FGSC16956 and FGSC12033, revealed unique nutritional phenotypes compared to the remaining set of 11 under-compensation mutants (Figure 3). FGSC16956 (ΔNCU02152) had the shortest period phenotype observed in the entire primary and secondary screens (Supplementary Table 1). NCU02152 was undescribed in the *Neurospora* literature but contains protein domain homology to the mammalian Cleavage & Polyadenylation Specificity Factor subunit 6 (*CPSF6*). CPSF6 is a member of the CFIm complex, which binds in the 3’ end of nascent mRNAs and facilitates cleavage and poly(A) tail placement. The CFIm complex also contains a second essential component CPSF5. Using the human CPSF5 protein sequence, NCU09014 was confidently identified as its *Neurospora* homolog (reciprocal BLAST e-values = 1e^-72^ / 3e^-74^). The single mutant FGSC12033 (ΔNCU09014 / Δ*cpsf5*) had the same short period length as Δ*cpsf6* (Supplementary Table 1), which indicates an obligate multimeric complex as with the mammalian CFIm complex. In the tertiary screen, Δ*cpsf5* and Δ*cpsf6* mutants showed progressive period shortening after ∼60 hours into the circadian free run, specifically when grown on high nutrient medium. Recalling the *prd-1* region-specific Nutritional Compensation phenotype where period defects were seen at the growth front encountering fresh medium, but not at the point of inoculation where nutrients were depleted (Emerson et al., 2015), we checked period length from old/aging tissue surrounding the point of fungal inoculation in Δ*cpsf5* and Δ*cpsf6* mutants. An ∼18-hour short period length was observed in the old tissue region of Δ*cpsf5* and Δ*cpsf6* mutants grown on high arginine (Figure 4A), compared to a ∼20-hour period length on zero nutrient medium (Supplementary Table 1). This result indicates that Δ*cpsf5* and Δ*cpsf6* mutants must undergo a transition from high-to-low amino acid levels in old tissue to reveal the 18-hour period defect (Supplementary Movie 2). The Δ*cpsf5* and Δ*cpsf6* mutants differ from other under-compensation mutants (Figure 3) because the short period defect was induced by amino acids, not by glucose, and because the Nutritional Compensation phenotype is specific to old tissue (Figure 4A).

**Figure 4.**
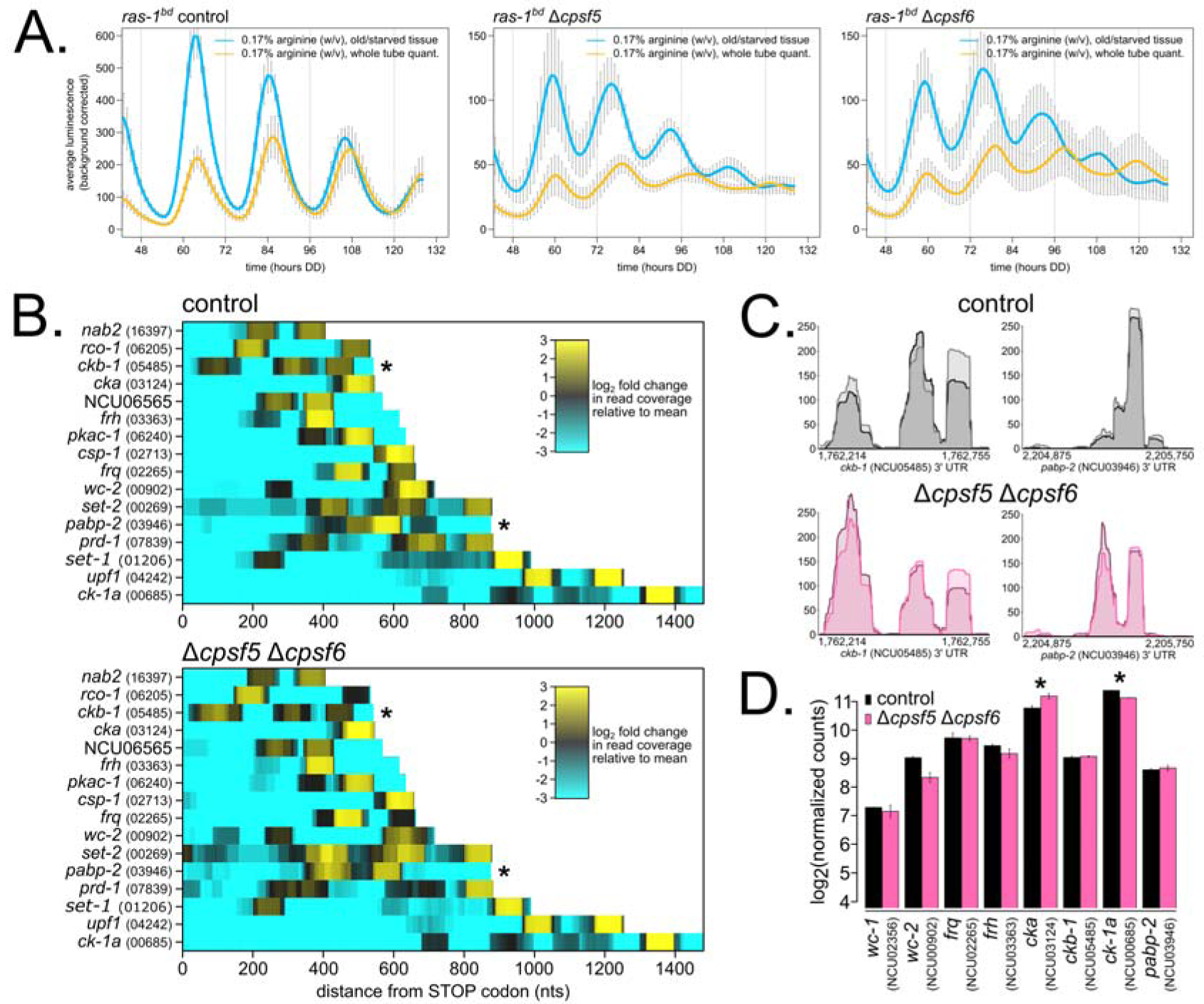
The *Neurospora* CFIm complex is involved in Alternative Polyadenylation (APA), and a subset of core clock and compensation-relevant genes are altered in ΔCFIm mutants. Circadian bioluminescence was recorded from race tube cultures of the indicated strains grown in high amino acid race tube medium (0.17% w/v arginine, 0% glucose). Luciferase signal was acquired from the entire race tube of fungal growth (yellow lines) or from an old tissue region of the race tube (blue lines) (see: Supplementary Movie 2). Circadian period lengths were computed for each region (Materials and Methods; N = 4 race tubes per genotype; standard deviation error bars): control: 21.7 ± 0.2 hrs (whole tube), 21.4 ± 0.3 hrs (old tissue); Δ*cpsf5*: 19.0 ± 0.5 hrs (whole tube), 17.2 ± 0.2 hrs (old tissue); Δ*cpsf6*: 19.4 ± 0.3 hrs (whole tube), 17.3 ± 0.3 hrs (old tissue) (**A**). Wild-type control and Δ*cpsf5* Δ*cpsf6* double mutant cellophane plate cultures were grown in constant light at 25°C for 72 hours on high nutrient medium (0.25% w/v glucose, 0.17% w/v arginine). Total RNA was extracted, and 3’ End Sequencing was performed (N = 2 biological replicates per genotype). 16 core clock and compensation genes of interest were selected, read pileup data from 3’ UTR regions were extracted, biological duplicate samples averaged together, and heatmaps were generated. Read count pileups are depicted as log_2_-fold changes, and both color scales are normalized to mean counts in the wild-type dataset. Each point along the x-axis represents nucleotide coordinates from the STOP codon for each mRNA (where data from any negative / Crick strand genes are shown in reverse orientation), and genes are ordered along the y-axis by increasing 3’ UTR lengths. Asterisks (*) indicate two genes, *ckb-1* and *pabp-2*, with significantly altered APA patterns between control and mutant (**B**). Genomic tracks were generated using the Gviz package in R to visualize poly(A) read pileups in the 3’ UTR regions of *ckb-1* and *pabp-2.* In the ΔCFIm mutant, poly(A) tail locations are significantly changed for *ckb-1* and *pabp-2* (**C**). Gene expression levels of core clock and compensation genes were measured by normalizing total read counts for each gene (Materials and Methods). Log_2_-transformed read counts are shown. Asterisks indicate p < 0.05 (*) by student’s t-test comparing mutant to control levels. The *cka^prd-3^* transcript is 1.34-fold upregulated, and the *ck-1a* transcript is 1.2-fold down-regulated in ΔCFIm (**D**).

### The Alternative Polyadenylation (APA) landscape is altered in Nutritional Compensation mutants

Given the large period defect observed in Δ*cpsf5* and Δ*cpsf6* mutants (Figure 4A) and other under-compensation mutants related to polyadenylation (Figure 3), we hypothesized that poly(A) tail maintenance and concurrently the stability of a core clock mRNA(s) is required for Nutritional Compensation. To assay the biochemistry of Δ*cpsf* mutants, we required a solid medium growth regime to carefully control nutrient levels and to confidently compare results to our genetic screen for Nutritional Compensation phenotypes. *Neurospora* biochemistry is traditionally accomplished using extracts from liquid-grown cultures (Nakashima, 1981; Kelliher et al., 2020b), where nutrient consumption rates are less well defined and less relevant to the ecological niche of the organism. A cellophane petri plate assay was developed to harvest biomolecules from solid medium cultures of *Neurospora* (Materials and Methods). We confirmed that the circadian clock is equally functional on cellophane plates and in liquid cultures (Supplementary Figure 5). We next generated a Δ*cpsf5* Δ*cpsf6* double mutant strain (hereafter referred to as ΔCFIm) and confirmed that its period length and Nutritional Compensation phenotype matched results from race tube assays (Supplementary Figure 5). Biological duplicate wild-type control and ΔCFIm mutant cellophane plates were grown at 25°C under constant light for 3 days on high nutrient medium (containing 0.25% w/v glucose 0.17% w/v arginine; high-to-low nutrient transition will occur in aged tissue before 72 hours growth), and total RNA was extracted for 3’ End Sequencing (Materials and Methods).

Nascent mRNA transcripts can contain multiple sites for polyadenylation to occur, which is known as Alternative Polyadenylation (APA). APA events generate mRNA isoforms with variable 3’ UTR lengths and nucleotide sequences (reviewed in: Mayr, 2017). Along with other factors involved in mRNA 3’ end cleavage and polyadenylation, CFIm mutants have been shown to alter APA patterns genome wide (Kubo et al., 2006; Martin et al., 2012; Zhu et al., 2018). Changes in APA have been directly linked to RNA stability in yeast and in mammals (Mayr and Bartel, 2009; Moqtaderi et al., 2022). We first sought to identify normal instances of APA in wild-type *Neurospora* 3’ UTRs by comparing our 3’ End Sequencing dataset to an existing 2P-Seq dataset (Zhou et al., 2018) (Materials and Methods). 843 consensus genes (>10% of *Neurospora* 3’ UTRs) contain multiple poly(A) sites after applying a strict intersection cutoff for identification in all 4 wild-type datasets (Supplementary Table 3), which is a lower genome-wide estimate than >60% APA in mammals. Next comparing ΔCFIm and controls, we found 193 examples of APA events in controls collapsing to a single poly(A) peak in mutants (21%), 123 examples of a single poly(A) peak in controls expanding to multiple APA events in mutants (13%), and 155 examples of APA events in both control and mutant where the location of the predominant poly(A) peak was significantly changed in mutants (16%) (Supplementary Table 4). Taken together, ∼50% of the APA landscape is altered in *Neurospora* ΔCFIm mutants. Knockdown of the mammalian CFIm complex causes global 3’ UTR shortening, as proximal poly(A) sites become preferred over distal poly(A) sites (Kubo et al., 2006; Martin et al., 2012; Zhu et al., 2018). In *Neurospora*, we find that a distal-to-proximal shift occurs in a majority (63%) of the 155 altered APA events in the ΔCFIm mutant.

Nine core clock and compensation genes are among the consensus list of 3’ UTRs with APA events: *ck-1a* (NCU00685), *frq* (NCU02265), *upf1^prd-6^* (NCU04242), *ckb-1* (*NCU05485*), *rco-1* (NCU06205), *pkac-1* (NCU06240), NCU06565, *prd-1* (NCU07839), and *nab2* (NCU16397) (Supplementary Table 3). Given the strict cutoff for consensus APA, we added 7 additional genes after visual inspection: *set-2* (NCU00269), *wc-2* (NCU00902), *set-1* (NCU01206), *csp-1* (NCU02713), *cka^prd-3^* (NCU03124), *frh* (NCU03363), and *pabp-2* (NCU03946). 3’ UTR regions containing APA events were visualized in a heatmap (Figure 4B). 2 out of 16 genes of interest, *ckb-1* and *pabp-2*, are among the list of genes with altered APA patterns in the ΔCFIm mutant (Supplementary Table 4; Figure 4B asterisks *). poly(A) read pileups were visualized for 3’ UTRs of *ckb-1* and *pabp-2* to confirm significant re-organization of poly(A) tail locations in the ΔCFIm mutant (Figure 4C). Following our original hypothesis, altered 3’ UTR length in ΔCFIm should lead to altered mRNA stability and changes in gene expression compared to control samples. An extremely slight and statistically non-significant increase (t-test, p = 0.5) was observed for both *ckb-1* and *pabp-2* gene expression levels (Figure 4D). *ckb-1* encodes the regulatory subunit of *Neurospora* Casein Kinase II (CKII), and interestingly, the catalytic alpha subunit of CKII (*cka^prd-3^*) increased significantly in the ΔCFIm mutant, despite no visible changes in 3’ UTR poly(A) sequence coverage for *cka^prd-3^* (Figure 4B). In addition, *ck-1a* gene expression levels are modestly (1.2 fold) but significantly decreased in ΔCFIm (Figure 4D), a result that is counterintuitive given the mutant’s short period length phenotype (Kelliher et al., 2020a). Future work will determine whether overexpression of CKII underlies the ΔCFIm short period length and Nutritional Compensation phenotypes.

### Chromatin modifiers emerge as key regulators of circadian period length across nutrients

We next examined the 4 over-compensation mutants identified in our genetic screen, where period lengthens as a function of nutrient levels (Figure 5A, Supplementary Table 1). The wild-type *Neurospora* clock is slightly under-compensated (Figures 1 – 3), and therefore this group of over-compensation mutants, together with *prd-1* (Figure 2B) (Emerson et al., 2015), represent clear and bona fide Nutritional Compensation defects. Protein Kinase A (*pkac-1*) shows extended compensation, or approximately the same period length at zero and high nutrients (Figure 5A). The effect of loss of PKA on Nutritional Compensation is subtle and likely due to its regulation of RCM-1 and WCC-independent *frq* transcription—RCM-1 normally acts as a general transcription co-repressor with RCO-1 and prevents *frq* transcription in a Δ*wc-1* or Δ*wc-2* background (Zhou et al., 2013; Liu et al., 2015); however, a Δ*rcm-1* mutant did have normal Nutritional Compensation in our screen (Supplementary Table 1). FGSC16412 (ΔNCU02961, Δ*rbg-28*) is a broadly conserved ribosome biogenesis factor in fungi, currently uncharacterized in the *Neurospora* literature but homologous to *RNH70* (YGR276C) in *S. cerevisiae*. Rnh70p (Rex1p) is a 3’-to-5’ exonuclease involved in 5S and 5.8S rRNA processing (Van Hoof et al., 2000). The completely arrhythmic phenotype of Δ*rbg-28* on high nutrients (Figure 5A) is intriguing and implicates translational machinery in Nutritional Compensation for the first time (see Discussion).

**Figure 5.**
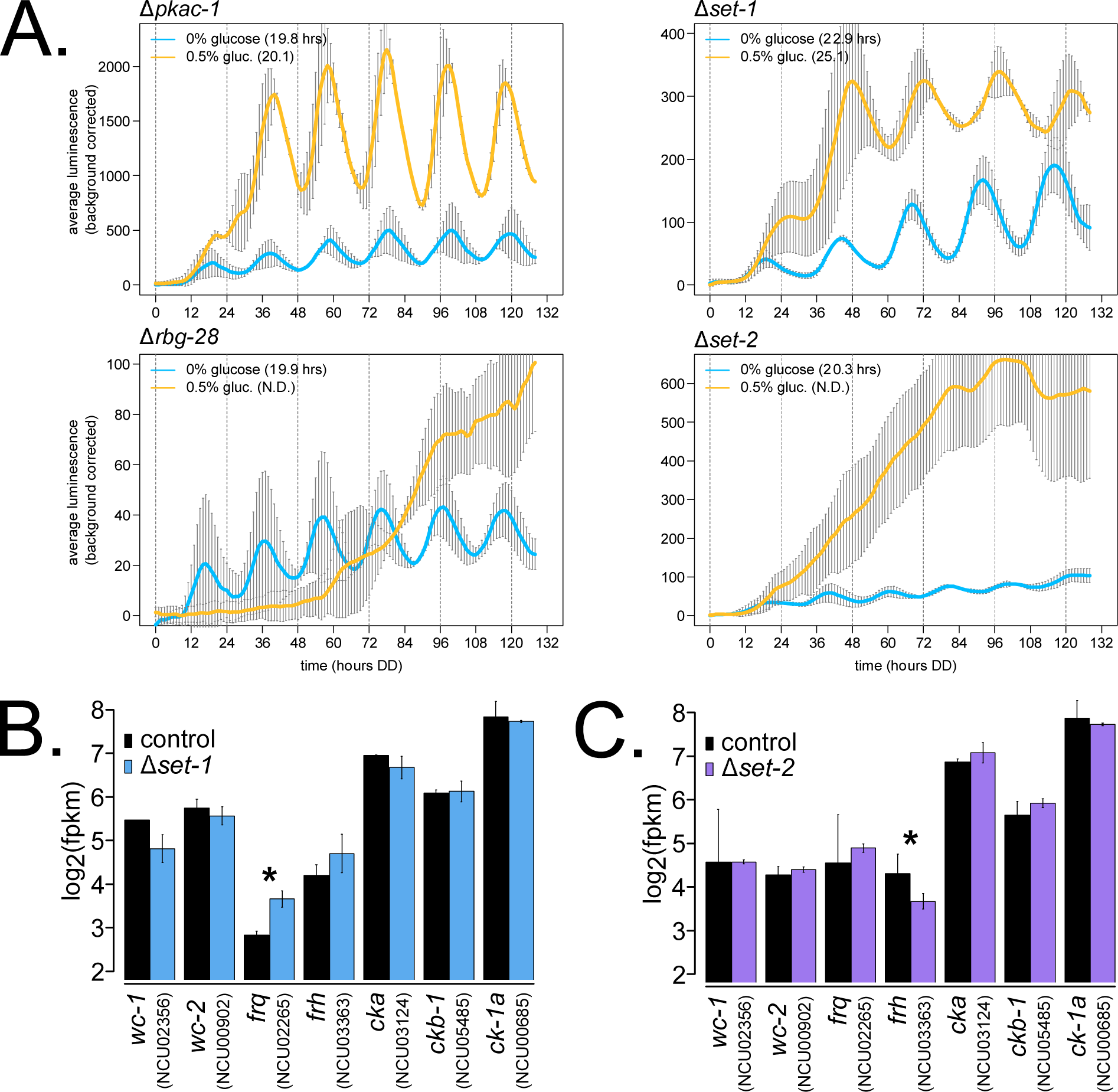
Nutritional over-compensation mutants are enriched for chromatin regulators. Circadian bioluminescence was recorded from race tube cultures of the indicated deletion mutants. High nutrient medium (yellow lines) contained 0.5% w/v glucose 0.17% w/v arginine, and zero nutrient medium (blue lines) contained 0% glucose 0% arginine. Period lengths are indicated as insets (N = 2 biological replicates per nutrient concentration) (**A**). RNA-Sequencing data were mined from a previous study (Zhu et al., 2019) (Materials and Methods), where 2% (high) glucose liquid cultures were harvested at circadian time point DD24. Log_2_-transformed FPKM values are shown for core clock gene expression levels (N = 2 biological replicates per genotype). The asterisk indicates p = 0.05 (*) by student’s t-test comparing mutant to control levels. The *frq* transcript is 1.78-fold more abundant in Δ*set-1* (**B**). RNA-Sequencing data were mined from a previous study (Bicocca et al., 2018) (Materials and Methods), where 1.5% (high) sucrose liquid cultures were sampled. Log_2_-transformed FPKM values are shown for core clock gene expression levels (N = 2 biological replicates for Δ*set-2*, and N = 6 biological replicates for controls). The asterisk indicates p < 0.05 (*) by student’s t-test comparing mutant to control levels. The *frh* transcript is 1.61-fold less abundant in Δ*set-2* (**C**).

Half of the novel over-compensation mutants are involved in chromatin regulation, and both contain SET domains (reviewed in: Freitag, 2017). SET-1 (NCU01206) is a histone H3 lysine 4 (K4) methyltransferase and the catalytic subunit of the COMPASS complex in *Neurospora*. Δ*set-1* was previously reported to be arrhythmic (Raduwan et al., 2013), but here we find an increasing period length as a function of glucose levels (Figure 5A). SET-1 was convincingly shown to regulate methylation levels and transcriptional repression at the *frq* locus (Raduwan et al., 2013; Zhu et al., 2019), and upon re-analysis of the Δ*set-1* transcriptome, *frq* is the only core clock gene significantly altered in the Δ*set-1* mutant due to loss of repression (Figure 5B). The Δ*set-1* nutritional over-compensation phenotype is consistent with loss of chromatin repression on *frq—*as glucose levels increase, more cellular energy is available for transcription/translation, and circadian period is lengthened due to high *frq* levels and prolonged negative feedback. This SET-1 nutritional mechanism is analogous to CKII’s role in regulating the larger pool of FRQ protein at higher temperatures in Temperature Compensation (Mehra et al., 2009).

SET-2 (NCU00269) is a histone H3 K36 methyltransferase in *Neurospora*, which deposits inhibitory chromatin marks in actively transcribed regions via a physical association with RNA Polymerase II and prevents improper transcription initiation inside coding regions (Adhvaryu et al., 2005; Bicocca et al., 2018). Δ*set-2* was also previously reported to be arrhythmic (Zhou et al., 2013; Sun et al., 2016), but here we find that Δ*set-2* rhythms are completely intact with a short period length on nutrient starvation medium (Figure 5A). SET-2 is required to maintain H3K36me2 and H3K36me3 marks across the *frq* locus, and the Δ mutant results in hyper-acetylation of *frq*, improper activation of WCC-independent *frq* transcription, constitutively high *frq* expression levels, and, presumably, the arrhythmic clock phenotype observed under high nutrients (Zhou et al., 2013; Sun et al., 2016). Our result indicates for the first time that WCC-independent *frq* transcription is nutrient dependent and only reaches levels sufficient for arrhythmicity at high nutrient levels. However, low WCC- independent *frq* transcription does not explain the short period length of the Δ*set*-2 mutant in zero nutrient medium (Figure 5A), as low WCC expression results in lengthened periods (Cheng et al., 2001). Curiously, upon mining existing data for the Δ*set-2* transcriptome, *frq* levels were not significantly increased, and instead *frh* levels were down-regulated (Figure 5C). *frh* is among the top down-regulated genes during glucose starvation (Supplementary Figure 4), and perhaps Δ*set-2* further affects *frh* transcription, leading to a circadian period change. Future work using cellophane plate assays (Supplementary Figure 5) will determine whether SET-2 regulation of *frh* or another gene expression program explains its short period length on nutrient starvation medium. Notably, two RNA helicases physically associated with the mammalian negative arm complex, DDX5 and DHX9, show a short period length upon siRNA knockdown (Padmanabhan et al., 2012), and so decreased levels of the *frh* helicase in Δ*set-2* could potentially explain its short period phenotype. Taken together, our genetic screen revealed mRNA stability, polyadenylation, and chromatin modifier pathways converging on gene expression regulation to enact circadian Nutritional Compensation in fungi.

### Nutritional Compensation also functions in a mammalian circadian system

Compensation is a defining principle of circadian oscillators. Temperature Compensation mechanisms are active in mammalian tissue culture (Izumo et al., 2003; Tsuchiya et al., 2003), despite long evolutionary timescales in homeothermic organisms. Thus, we hypothesized that Nutritional Compensation is also a conserved feature between the fungal and mammalian circadian clocks. Four previous studies hinted that Nutritional Compensation mechanisms may be actively maintaining circadian period length across physiologically relevant nutrient environments. Inhibiting transcription using α-amanitin or actinomycin D against RNA Polymerase II activity led to dose-dependent shortening of the NIH3T3 period length (Dibner et al., 2009). In other words, the mammalian circadian oscillator is over-compensated with respect to transcription rates, reminiscent of temperature over-compensation observed in *Gonyaulax* (Hastings and Sweeney, 1957). Similarly, induction of autophagy and amino acid starvation shortens period length in MEFs (Beesley et al., 2020), once again indicating an over-compensation phenotype to amino acid levels in mammals. Varying levels of FBS do not alter the circadian period length in NIH3T3 cells (Matsumura et al., 2014) despite substantial changes in cell growth rate. Rhythms at the single-cell level of MEFs grown in microfluidic devices show slight period shortening when cells were given fresh medium every hour compared to a single medium supply at the beginning of the experiment, suggesting instead nutritional under-compensation (Gagliano et al., 2021). Thus, preliminary evidence is consistent with functional Nutritional Compensation in the mammalian circadian system.

We used a U2OS *Bmal1*-*dLuc* reporter cell line to compare rhythms in high versus low glucose medium and found a slight under-compensation phenotype (Supplementary Figure 6), similar to the single-cell rhythms report (Gagliano et al., 2021). Period length was 0.5 hours shorter in 25 mM (high) glucose compared to 5.56 mM (low) glucose, trending in the same direction as the ∼1.5-hour period shortening seen for the wild-type *Neurospora* clock across glucose concentrations (Figures 1 – 3). In this circadian assay, *Bmal1*-*dLuc* cells have reached confluency and are no longer actively dividing, but metabolism and drug responses of cultured cells are reported to be significantly different in high versus low glucose tissue culture medium (reviewed in: Abbas et al., 2021).

Maintenance of circadian period length between high and low glucose medium and a small amount of literature precedent does not, however, prove the relevance of Nutritional Compensation to the mammalian clock. A mutant phenotype showing nutrient-dependent period changes (i.e. defective Nutritional Compensation) would provide much stronger evidence for the biological relevance of compensation. Therefore, we selected two of the most significant compensation phenotypes from our fungal genetic screen—Δ*cpsf6* (NCU02152) and Δ*set-2* (NCU00269)—and identified the homologous human genes *CPSF6* and *SETD2* (reciprocal BLAST e-values = 7e^-6^ / 7e^-8^ for Cpsf6 and 4.4e^-61^ / 5.7e^-62^ for Set-2). If the circadian functions of *CPSF6* or *SETD2* are indeed conserved with *Neurospora*, we expected to observe a period length change as well as a Nutritional Compensation defect. *CPSF6* and *SETD2* were not among the hits from a genome-wide screen for period length defects using U2OS cells (Zhang et al., 2009) or tested in a kinase/phosphatase siRNA screen (Maier et al., 2009); however, visual inspection of the genome-wide screen data confirmed that a subset of the pooled siRNAs did show period effects after *CPSF6* or *SETD2* knockdown (source: BioGPS database, “Circadian Genomics Screen” plugin).

*CPSF6* and *SETD2* were knocked down using siRNAs in *Bmal1*-*dLuc* cells, and period length was measured from high and low glucose medium (Figure 6). AllStars Negative Control siRNA, which does not target any known mammalian transcript, was used as an internal control for each biological replicate assay, and *CRY2* siRNA knockdown was used as a positive control for a known long period phenotype (Baggs et al., 2009; Lee et al., 2019). Control *Bmal1*-*dLuc* cells had a slight under-compensation phenotype (Figure 6A), matching preliminary results (Supplementary Figure 6). *CRY2* knockdown lengthened period by ∼4 hours in both high and low glucose conditions, indicating no effect on Nutritional Compensation (Figure 6A). *CPSF6* knockdown lengthened period by ∼1.5 hours in high glucose and further lengthened period by ∼3 hours in low glucose, indicating an under-compensation phenotype compared to control cells (Figure 6A). The long period observed in *CPSF6* knockdowns was the opposite of the short period phenotype in *Neurospora* (Figure 4A). Most interestingly, *SETD2* knockdown drastically increased the amplitude of the *Bmal1*-*dLuc* transcriptional reporter compared to controls. As with *Neurospora*, *SETD2* knockdown rhythms were less robust (Figures 5A & 6A). *SETD2* knockdown lengthened period by ∼2.5 hours in high glucose but only lengthened period by ∼1 hour in low glucose (Figure 6A). To further validate nutritional over-compensation in *SETD2* knockdowns, *Bmal1*-*dLuc* cells were compared across higher concentrations of glucose with more robust rhythms (Figure 6B). Just like *Neurospora*, *SETD2* mutants show a nutritional over- compensation phenotype. This genetic evidence strongly suggests that Nutritional Compensation mechanisms also regulate the mammalian circadian clock in physiologically relevant environments.

**Figure 6.**
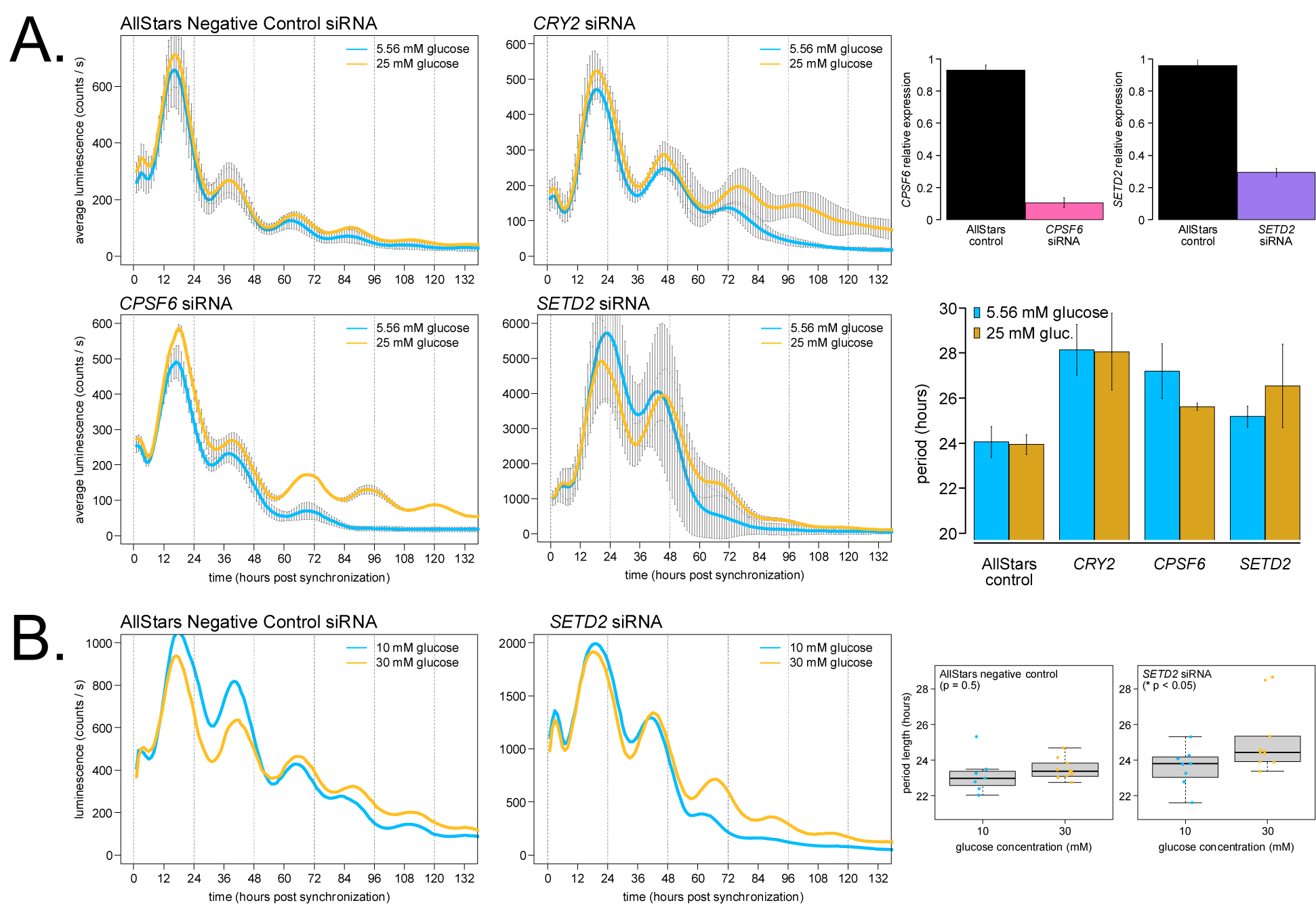
*CPSF6* and *SETD2* are required for normal Nutritional Compensation of U2OS *Bmal1-dLuc* cells cultured in high versus low glucose. U2OS cells were transfected with the indicated siRNAs (15 pmol total), incubated for 2 days, and *Bmal1-dLuc* rhythms were measured for 5 – 6 days following dexamethasone synchronization. Knocked-down cells were assayed in either MEM high glucose (yellow lines) or MEM low glucose (blue lines) medium. Averaged luciferase traces are shown for each glucose concentration without detrending or further data manipulation (N = 6 biological / technical replicates for AllStars negative controls; N = 4 biological / technical replicates for other siRNA knockdowns per nutrient concentration; standard deviation error bars). siRNA knockdown efficiency was measured using RT-qPCR relative to non-transfected control samples (Materials and Methods). ∼90% knockdown was achieved for *CPSF6*, and ∼70% knockdown for *SETD2* under these experimental conditions (N = 2 biological replicate samples each with N = 3 technical replicate RT-qPCR reactions). Period lengths were calculated and averaged for all replicates (**A**). siRNA knocked-down cells were assayed in either DMEM 30 mM glucose (yellow lines) or DMEM 10 mM glucose (blue lines) medium. One representative luciferase trace is shown (N = 7 – 10 biological / technical replicates per nutrient concentration). Period lengths were calculated and averaged for all replicates and plotted as boxplots (**B**). Average period lengths were: 23.2 ± 1.1 hrs (low glucose control) and 23.5 ± 0.6 hrs (high glucose control); 23.6 ± 1.1 hrs (low glucose *SETD2* knockdown) and 25.2 ± 1.8 hrs (high glucose *SETD2* knockdown) (* p < 0.05, student’s t-test).

## Discussion

We present the largest genetic screen to date for mutants displaying altered circadian Nutritional Compensation (Supplementary Tables 1 & 2, Figure 2C). Together with a recent survey of 177 transcription factor knockouts (Muñoz-Guzmán et al., 2021), circadian functional genomics is well underway utilizing the *Neurospora* deletion collection (Colot et al., 2006). In this study, 16 new Nutritional Compensation mutants were identified and characterized along with the 4 previously characterized mutants (Δ*csp-1*, *prd-1*, Δ*rco-1*, and Δ*ras2*)—fungal Nutritional Compensation occurs at the level of gene regulation and involves transcription factors, RNA helicases, chromatin modifiers, and polyadenylation machinery. Nutritional Compensation effectors are responsible for directly regulating the core oscillator to maintain the circadian period length across different nutrient levels. In high nutrient environments, CSP-1 forms a negative feedback loop on WCC expression and activity by inhibiting *wc-1* overexpression and preventing nutritional under-compensation (Sancar et al., 2012; Dovzhenok et al., 2015). RCO-1, SET-2, and possibly PKA are required to block WCC-independent *frq* transcription in high nutrient conditions (Figure 5) (Zhou et al., 2013; Olivares-Yanez et al., 2016; Sun et al., 2016). More broadly, repressive chromatin marks at the *frq* locus, including those regulated by SET-2, SET-1, and the COMPASS complex, are required to maintain Nutritional Compensation, especially in high nutrients (Figure 5). The nonsense-mediated decay machinery is required for circadian function through its regulation of *ck-1a* expression (Kelliher et al., 2020a), and *ck-1a* transcript stability appears to undergo additional, nutrient-dependent regulation because deletion of the *ck-1a* 3’ UTR (its NMD-targeting sequence) causes nutritional under-compensation (Figure 3, Supplementary Figure 3). The alternative polyadenylation (APA) landscape in *Neurospora* is dynamic across nutrient conditions and requires CFIm complex activity (Figure 4).

This screen and associated candidates have opened up several avenues for further work on Nutritional Compensation mechanisms in *Neurospora*. For instance, PRD-1 RNA helicase target genes under high glucose conditions are completely unknown. In this study, we provide some evidence that CKII activity is altered in ΔCFIm mutants (Figure 4), but the effect of CKII overexpression on both circadian period length and Nutritional Compensation remains to be tested. Genetic screens extending from this work can examine both transcription factor and chromatin modifier knockouts for Nutritional Compensation phenotypes. Curiously, the subset of chromatin readers and writers that have been screened to date (using standard medium conditions) have an unusually high ∼15% hit rate for circadian period defects (Belden et al., 2007b; Wang et al., 2014). In this study, chromatin modifiers not only regulate chromatin state at the *frq* locus, but dynamic expression of *frh* across nutrient levels may also be involved in compensation, potentially mediated by SET-2 (Figure 5C). Intriguingly, quantitative modeling has suggested that nutrient-dependent FRH sequestration away from the negative arm complex is a plausible mechanism for Nutritional Compensation (Upadhyay et al., 2019). Mechanistic work in prokaryotes has indicated that Nutritional Compensation can be derived from the same core clock enzyme, KaiC, through its protein domains with equal and opposite balancing reaction rates (Phong et al., 2013; Hong et al., 2020). In fungi and perhaps other eukaryotic clocks, Nutritional Compensation appears to be maintained through regulation of multiple different core clock factors. Nutritional Compensation mechanisms very likely extend to other physiological environmental variables such as nitrogen sources and levels (Huberman et al., 2021), vitamins, and soil pH (Ruoff et al., 2000).

The work establishes Nutritional Compensation in the mammalian circadian clock for the first time, and its clinical relevance may be high. There is a direct link between pancreatic clock function and risk for type 2 diabetes (Marcheva et al., 2010). Fasting glucose concentration in human serum is approximately 5 mM in healthy controls, but glucose levels can increase by 3+ orders of magnitude in type 2 diabetic hyperglycemia (Shapiro et al., 1991; Radziuk and Pye, 2006). Thus, deleterious SNPs in Nutritional Compensation-relevant genes could exacerbate disease outcomes when cellular clocks encounter nutrients outside of the physiological range. In addition to Nutritional Compensation, there is a large body of quality literature describing the set of metabolites and metabolic enzymes that can directly feed back and affect circadian function, including adenosine monophosphate (AMP) and AMP kinase, nicotinamide adenine dinucleotide (NAD+) and SIRT1 activity, acetylglucosamine (O-GlcNAc) (Liu et al., 2021), and mTOR activity (Ramanathan et al., 2018) (reviewed in: Bass and Takahashi, 2010; Sancar and Brunner, 2014; Asher and Sassone-Corsi, 2015; Dibner and Schibler, 2015). In *Neurospora*, extensive metabolic rhythms are also present (Hurley et al., 2018; Baek et al., 2019), and rhythmic metabolic reaction fluxes likely function similarly to the mammalian clock (Krishnaiah et al., 2017; Thurley et al., 2017; Collins et al., 2021). For the mammalian clock, it will be critical to define “physiologically relevant” nutrient levels, which could vary by organ or even cell type. A high fat diet has been shown to lengthen circadian behavioral rhythms by ∼0.5 hours (Kohsaka et al., 2007), suggesting an over-compensation phenotype. In this study, we observed nutritional under-compensation in U2OS osteosarcoma cells (Figure 6). Does a high fat diet push cells out of the physiological range of compensation for the circadian oscillator, or are different mammalian cell types differentially over- or under-compensated to nutrients? Notably, time-restricted feeding has been shown to alleviate circadian disruption and some metabolic consequences of a high fat diet (Hatori et al., 2012; Wehrens et al., 2017). Future work will implement more metabolically relevant cell types, such as hepatocytes and adipocytes (Ramanathan et al., 2014), to answer these questions about mammalian clock compensation.

Circadian control of polyadenylation is an emerging topic. Modern transcriptomic approaches have identified hundreds of rhythmic Alternative Polyadenylation (APA) events in mice and in plants (Liu et al., 2013; Gendreau et al., 2018; Greenwell et al., 2020; Yang et al., 2020). We and others have begun to define the set of APA events in *Neurospora* (Zhou et al., 2018) (Supplementary Table 3), which can be extended to circadian APA events. Two cleavage and polyadenylation factors (CPSF1 and 7) were found to physically interact with the negative arm complex in mice (Ju et al., 2020), and *CPSF6* oscillates at the transcriptional level in mouse kidney and brain (Zhang et al., 2014). During preparation of this manuscript, circadian colleagues reported the long period length of *CPSF6* knockdown (Schmal et al., 2021), which our results further support (Figure 6A). Interestingly, *CPSF6* knockdown cells showed a temperature under-compensation defect, and multi-omics identified *EIF2S1* as the key effector gene upon *CPSF6* knockdown (Schmal et al., 2021). The *Neurospora* homolog of *EIF2S1* is eIF2α / NCU08277 (reciprocal BLAST e-values = 6^e-91^ / 1e^-81^), which is a central hub of rhythmic translation initiation peaking in the subjective evening (Karki et al., 2020). We identified a ribosome biogenesis exonuclease, RBG-28, which is required for rhythmicity under high nutrient conditions (Figure 5). Transcription, RNA processing reactions (such as nonsense-mediated decay, splicing, and polyadenylation), and translation are tightly coupled processes. In fact, rhythmic polyadenylation of rRNAs has been linked to translational rhythms in mouse liver (Sinturel et al., 2017). Nutritional Compensation pathways likely occur at multiple steps in the gene expression of multiple core clock components.

By suggesting *CPSF6* and *SETD2* as targets, *Neurospora* functional genetics has again informed mammalian-relevant circadian mechanisms (reviewed in: Loros, 2020), and the over- compensation defect of *SETD2* provides affirming evidence of eukaryotic Nutritional Compensation outside of *Neurospora* (Figure 6). A handful of previous studies have implicated histone methyltransferases in mammalian circadian function, including both activating (Katada and Sassone-Corsi, 2010; Valekunja et al., 2013) and repressive chromatin marks (Etchegaray et al., 2006). *SETD2* joins a growing list of circadianly-relevant histone methyltransferases and chromatin modifiers. In fact, recent work has demonstrated a novel function for a key histone methyltransferase in the circadian transcription-translation feedback loop (TRITHORAX in insect, MLL1 in mammals), further highlighting the utility of circadian model systems for understanding the mammalian clock (Zhang et al., 2022).

## Materials and Methods

### *Neurospora* strains, growth conditions, and genetic screen

Strains used in this study were derived from the wild-type background (FGSC2489 *mat* A), *ras-1^bd^* background (87-3 *mat* a or 328-4 *mat* A), or the Fungal Genetics Stock Center (FGSC) knockout collection as indicated (Supplementary Table 1). Strains were constructed by transformation or by sexual crosses using standard *Neurospora* methods (http://www.fgsc.net/Neurospora/NeurosporaProtocolGuide.htm). The *frq* clock box transcriptional reporter was transformed and used as previously described (Kelliher et al., 2020a). The fungal biomass reporter gene (Supplementary Figure 1) is composed of 430 bp of the *gpd* promoter from *Cochliobolus heterostrophus* driving constitutive levels of codon optimized *luciferase* and integrated at the *csr-1* (NCU00726) locus (Bartholomai, 2021).

All race tubes contain a base medium of 1X Vogel’s Salts, 1.5% w/v Noble agar (Thermo Fisher # J10907), and 50 ng/ml biotin. Noble agar was used instead of standard bacteriological agar (Thermo Fisher # J10906) because impurities in bacteriological agar can be metabolized by *Neurospora* and interfere with accurate quantification of Nutritional Compensation phenotypes (Emerson et al., 2015). Glucose and arginine were supplemented into race tube medium as indicated. High glucose was defined as 0.5% w/v (27.8 mM) based on literature precedent (Sancar et al., 2012; Olivares-Yanez et al., 2016) and based on growth rate and period length similarity to higher glucose concentrations (Figure 1A, Supplementary Figure 1G). High arginine was defined as 0.17% w/v because concentrations higher than 0.5% w/v interfere with circadian banding in race tube assays (Sargent and Kaltenborn, 1972). To optimally visualize and quantify the circadian banding pattern in the primary genetic screen (Figure 2C), screen medium contained: 1X Vogel’s Salts, 1.5% Noble agar, 50 ng/ml biotin, and 0.17% arginine. To quantify period lengths in carbon and nitrogen starvation conditions, the secondary screen medium contained: 1X Vogel’s Salts, 1.5% Noble agar, 50 ng/ml biotin, and 25 μM luciferin (GoldBio # 115144-35-9). The tertiary screen medium contained: 1X Vogel’s Salts, 1.5% Noble agar, 50 ng/ml biotin, 25 μM luciferin, and glucose/arginine levels as indicated (Figures 3 – 5). For Temperature Compensation 96-well plate experiments (Figure 2A), the standard medium recipe contained: 1X Vogel’s Salts, 1.5% bacteriological agar, 50 ng/ml biotin, 25 μM luciferin, 0.03% w/v glucose, and 0.05% w/v arginine.

Liquid medium cultures were grown from fungal plugs in Bird Medium + 1.8% w/v glucose (Supplementary Figure 5) as previously described (Kelliher et al., 2020b). Solid medium cultures were implemented to determine mRNA (or protein) levels from Nutritional Compensation mutants of interest. Medium was poured into 100-mm petri plates and cooled to solidify (∼20 ml per plate; 1X Vogel’s Salts, 1.5% Noble agar, 50 ng/ml biotin, 25 μM luciferin, and glucose/arginine levels as indicated). Cellophane (Idea Scientific # 1080) paper discs were cut to the size of 100-mm plates, and autoclaved to sterilize. A sterile cellophane disc was then placed on top of the solidified medium. Conidia from strains of interest were resuspended in 100 μl of sterile water, vortexed to mix, pipetted on top of the cellophane disc, and spread with a sterile plate spreader (in order to maintain approximately the same conidial age across the cellophane plate). Inoculated plates were then covered with a Breathe-Easy strip for gas exchange (USA Scientific # 9123-6100). After growing tissue on cellophane plates for the indicated amount of time, mycelia and conidia were harvested from atop the cellophane layer by scraping with a 1000 μl pipette tip. Harvested fungal tissue (approximately the size of one US quarter per each 100-mm cellophane plate) was rapidly hand dried using paper towels and an Eppendorf tube rack, and flash frozen in liquid nitrogen for storage before biochemical extraction.

Most strains were genotyped by growth on selective medium (5 μg/ml cyclosporine A and/or 200-300 μg/ml Hygromycin). Key strains were genotyped by PCR as previously described (Kelliher et al., 2020a) using genotyping primers: *ck-1a^LONG^-VHF*Δ*3’UTR*::hyg^R^ (NCU00685 Δ3’UTR): 5’ GCTGCTGCTCGTAAGGAC 3’ and 5’ CATCAGCTCATCGAGAGCCTG 3’

Δ*cpsf5*::hyg^R^ (FGSC KO mutant): 5’ CTCTGGTCGAGAACACTGCG 3’ and 5’ CAGGCTCTCGATGAGCTGATG 3’

Δ*cpsf6*::hyg^R^ (FGSC KO mutant): 5’ CACCAACCCTAACCCGTGAT 3’ and 5’ CAGGCTCTCGATGAGCTGATG 3’

Δ*set-2*::hyg^R^ (FGSC KO mutant): 5’ GACGTCATCGGTGTTGAGAC 3’ and 5’ CAGGCTCTCGATGAGCTGATG 3’

### *Neurospora* luciferase reporter detection and data analysis

96-well plates were inoculated with conidial suspensions and entrained in 12 hour light:dark cycles for 2 days in a Percival incubator at 25°C. Temperature inside the Percival incubator was monitored using a HOBO logger device (Onset # MX2202) during entrainment and free run. Race tubes were entrained in constant light at 25°C for 3 – 24 hours (mean entrainment time for all experiments was 14 hours in LL / overnight). Entrained 96-well plates or race tubes were then transferred into constant darkness to initiate the circadian free run. Individual race tubes were separated by ∼3 cm tall strips of 6-ply black railroad board paper to prevent contamination of light signal between cultures. Luminescence was recorded using a Pixis 1024B CCD camera (Princeton Instruments). Bioluminescent signal was acquired for 10 –15 minutes every hour using LightField software (Princeton Instruments, 64-bit version 6.10.1).

The average bioluminescent intensity of each 96-well or race tube was determined using a custom ImageJ Macro with background correction for each image (Larrondo et al., 2012, 2015). Most race tube period lengths reported in this study were derived from luciferase signal measurements across the entire race tube (Figures 1, 3, and 5). However, the long period defect in the *prd-1* mutant only occurs at the growth front (i.e. high nutrients) region of fungal tissue (Emerson et al., 2015). For the *prd-1* strain, an ImageJ macro was modified to quantify only the fungal growth front (Figure 2B). On the other hand, the Δ*cpsf5* and Δ*cpsf6* mutants showed additional period shortening in aged tissue (Figure 4A) (Supplementary Movie 2). For the *cpsf* mutants, an ImageJ macro was modified to quantify only the old tissue region. Custom ImageJ Macros to quantify the growth front or old tissue regions of race tubes from Princeton *.spe image files are available at: https://github.com/cmk35. To calculate the circadian period length, background-corrected luminescence traces were run through two different algorithms and averaged as previously described (Kelliher et al 2020 eLife). Race tubes period lengths were measured using ChronOSX 2.1 software. For Temperature Compensation experiments, the Q_10_ temperature coefficient was calculated using the formula: [(frequency of clock at 30°C) / (frequency of clock at 20°C)] ^[10°C / (30°C - 20°C)]^, where frequency = period length^-1^.

### *Neurospora* RNA isolation and 3’ End Sequencing analyses

Frozen *Neurospora* tissue was ground in liquid nitrogen with a mortar and pestle. Total RNA was extracted with TRIzol (Invitrogen # 15596026) and the Direct-zol RNA MicroPrep kit (Zymo Research # R2060) according to the manufacturer’s instructions and including the on- column DNAse I treatment step (Roche # 04 716 728 001, 10 U/μl stock, 30 U used per sample). Total RNA samples were prepared for Northern Blotting, 3’ End Sequencing, or stored at -80°C.

Northern blotting was performed as previously described (Kelliher et al., 2020a) with slight modifications. Equal amounts of total RNA (7 μg) were loaded per lane of a 0.8% w/v agarose gel (Supplementary Figure 5). For blot visualization, anti-Digoxigenin-AP Fab fragments was purchased from Sigma (Roche # 11 093 274 910) and used at 1:10,000 (75 mU/ml).

Total RNA was submitted to the Dartmouth Genomics Shared Resource (GSR) for 3’ end library preparation and sequencing. 75 bp single-end (SE) strand-specific libraries were prepared using the Lexogen QuantSeq 3’ REV kit, multiplexed, and sequenced on an Illumina Mini-Seq. 6.92 ± 0.30 million reads were obtained for each sample, and read quality was confirmed using FastQC. Raw FASTQ files were aligned to the *Neurospora crassa* OR74A NC12 genome (FungiDB version 45 accessed October 25, 2019) using STAR (Dobin et al., 2013). 91.5 – 93.5% of the reads mapped uniquely to the NC12 genome. Because 3’ end libraries generate only 1 sequencing read at the extreme 3’ end of a given mRNA transcript (directly before its poly(A) tail), gene expression was quantified by counting reads assigned to each genetic locus using HTSeq-count (Anders et al., 2015). Gene count normalization by library size between samples was performed using a custom R script. 3’ End Sequencing data have been submitted to the NCBI Gene Expression Omnibus (GEO; https://www.ncbi.nlm.nih.gov/geo/) under accession number GSE201901.

RNA-Sequencing datasets from 4 other studies were mined in the analyses presented. To examine wild-type *Neurospora* gene expression under carbon starvation (Supplementary Figure 4, Supplementary Table 2), RNA-seq data were taken from a study where liquid cultures (25°C, LL) were grown for 16 hours in 1X Vogel’s 2% sucrose minimal medium and shifted to either 0% or 2% glucose 1X Vogel’s medium for 60 minutes (Wang et al., 2017) (GSE78952). To examine gene expression in the Δ*set-1* mutant background (Figure 5B), RNA-seq data were taken from a study where liquid cultures (25°C, DD24) were grown in 2% glucose Liquid Culture Medium for 48 hours of total growth (Zhu et al., 2019) (GSE121356). To examine gene expression in the Δ*set-2* mutant (Figure 5C), RNA-seq data were taken from a study where liquid cultures (32°C) were grown in 1X Vogel’s 1.5% sucrose medium (Bicocca et al., 2018) (GSE82222 and GSE118495). These three RNA-Seq datasets were re-processed exactly as previously described (Kelliher et al., 2020a), and FPKM gene expression values were used in the analyses presented. To examine and compare wild-type poly(A) tail locations (Supplementary Table 3), 2P-Seq data were taken from a previous study where nuclear fractions were isolated from 1X Vogel’s 2% glucose liquid medium (Zhou et al., 2018) (SRA PRJNA419320). Raw 2P-Seq data were filtered according to custom a Perl script from the original study (“Step 1”: https://github.com/elifesciences-publications/poly-A-seq). After read filtering, duplicate 2P-Seq FASTQ files were processed in exactly the same manner as the new 3’ End Sequencing dataset generated in this study.

NC12 mapped reads from 3’ End Sequencing (this study) and 2P-Seq (Zhou et al., 2018) data were sorted and indexed using Samtools (BAM file outputs) and then visualized using IGV. Read pileups denoted the location of poly(A) tails in both datasets. To map locations of poly(A) tails genome wide, the ChIP-Seq peak calling algorithm MACS2 was re-purposed (Zhang et al., 2008). The relevant MACS2 parameters used to identify poly(A) peaks were: effective genome size (-g 4.014e7), retention of duplicate reads in pileups (--keep-dup all), summit and subpeak identification (--call-summits), fragment size estimation and shifting turned off (--nomodel --extsize 75), and a false discovery rate cutoff for significant peaks (-q 0.01). MACS2 peaks were assigned to the corresponding gene 3’ UTR region using a custom R script. The *Neurospora crassa* NC12 transcriptome annotation remains partially incomplete with only 7,793 out of 10,591 unique NCU IDs having 3’ UTRs annotated. As a result, ∼14-18% of all MACS2 peaks were pruned from consideration due to missing annotations. Importantly, there are also examples of under-annotated 3’ UTR regions, where the poly(A) read pileup signal is clearly located outside of the 3’ UTR annotation. One such critical example occurs at the *frequency* locus (positive / Watson strand gene), where the predominant poly(A) peak is centered at LG VII coordinate 3,136,633, and its longest 3’ UTR annotation ends at coordinate 3,136,464. The *Neurospora* NC12 transcriptome annotation was last updated in March 2015 before migration from the Broad Institute to the FungiDB database (Basenko et al., 2018). The 3’ End Sequencing analyses presented here can be updated upon release of an improved transcriptome annotation. Furthermore, there are 621 instances of overlapping coordinates within the 3’ UTRs of tail-to-tail oriented genes, and any poly(A) peaks falling in these gene assignment ambiguous regions were also removed from consideration (∼9% of MACS2 peaks). The remaining MACS2 peaks (∼6,600 unique poly(A) peaks per sample) were assigned to the corresponding gene 3’ UTR region and analyzed using custom R scripts. Alternative Polyadenylation (APA) events were defined as instances of more than one distinct MACS2 peak assigned to a single 3’ UTR region. poly(A) tail read pileups from genes of interest were extracted using the igvtools count function, and genome coordinate plots were generated using the R Bioconductor package Gviz (Figure 4C). 3’ UTR heatmaps were generated using a custom R script (Figure 4B). All custom R scripts for gene expression analyses and alternative polyadenylation analyses are available at: https://github.com/cmk35.

### Mammalian cell culture, synchronization, and siRNA knockdown reagents

U2OS cells were stably transfected under puromycin selection using a construct containing the mouse *BMAL1* promoter (Sato et al., 2006; Gamsby et al., 2009) driving destabilized luciferase (Ueda et al., 2002). U2OS-m*BMAL1*-*dLuc*-Puro (referred to as: *Bmal1*- *dLuc*) cells were maintained at 37°C and 6% CO_2_ in 25 mM (high) glucose DMEM (Thermo Fisher # 11995-065 with 1 mM sodium pyruvate; or Thermo Fisher # 11965-092 without pyruvate) supplemented with 10% v/v FBS (Thermo Fisher # 10437-036, LOT # 2199672RP) and 1.5 μg/ml of puromycin (Sigma # P9620, 10 mg/ml stock).

For control Nutritional Compensation assays (Supplementary Figure 6), *Bmal1*-*dLuc* cells were subcultured from the same 100-mm dish and grown to 95-100% confluence in 35-mm dishes (Corning # 430165) containing 2 ml of DMEM 25 mM glucose, 10% FBS, and puromycin. Confluent cells were washed once in warm 1X PBS pH 7.4 (Corning # 21-040-CV). The medium was changed to either 2 ml of DMEM 25 mM high glucose (Thermo Fisher # 11995-065 with 1 mM sodium pyruvate) or 2 ml of DMEM 5.56 mM low glucose (Thermo Fisher # 11885-084 with 1 mM sodium pyruvate). Both synchronization-release medium formulations were pre-warmed and each contained: 10% v/v FBS, 1.5 μg/ml puromycin, 0.1 mM luciferin (GoldBio, 0.1 M stock), and 0.1 μM dexamethasone (Sigma # D2915, 1 mM stock). Dexamethasone is used to reset cells to the same circadian phase and initiate the circadian free run for recording.

For siRNA knockdown assays (Figure 6), *Bmal1*-*dLuc* cells were subcultured from the same 100-mm dish and grown to 60-80% confluence in 35-mm dishes containing DMEM 25 mM glucose, 10% FBS, and puromycin. Cells were washed once in 1X PBS, and the medium was changed to 2 ml of Opti-MEM (Thermo Fisher # 31985-070) with 5% v/v FBS. Cells were transfected with the indicated siRNAs (15 pmol of total siRNA per 35-mm dish) (Baggs et al., 2009) using the Lipofectamine 3000 transfection reagent (Thermo Fisher # L3000) and according to the manufacturer’s instructions for 6-well plates. Although the Opti-MEM medium formulation is not publicly available, one study reported the Opti-MEM glucose concentration as 2.5 g/L or 13.88 mM (Young et al., 2004). siRNAs were obtained from Qiagen: AllStars Negative Control siRNA (Qiagen # 1027280); human *SETD2* siRNA (Qiagen # 1027416, FlexTube GeneSolution GS29072, 4x siRNAs used at 3.75 pmol each); human *CPSF6* siRNA (Qiagen # 1027416, FlexTube GeneSolution GS11052, 4x siRNAs used at 3.75 pmol each); human *CRY2* siRNA (Qiagen # 1027416, FlexTube GeneSolution GS1408, 4x siRNAs used at 3.75 pmol each) (Lee et al., 2019). Cells were incubated for 2 days before removing the siRNA transfection medium and proceeding with RNA extraction or with circadian recordings.

RT-qPCR was used to validate siRNA knockdown efficiencies. 2-day transfected cells were washed once in 1 ml of ice-cold 1X PBS. Cells were harvested by scraping in 1 ml of TRIzol (Invitrogen), and total RNA extraction was performed according to the manufacturer’s instructions. 500 ng of mRNA was converted into cDNA using the oligo(dT) method from the SuperScript IV First-Strand synthesis kit (Invitrogen # 18091–050). RT-qPCR was performed using SYBR green master mix (Qiagen # 204054) and a StepOne Plus Real-Time PCR System (Applied Biosystems). C_t_ values were determined using StepOne software (Life Technologies) and normalized to the *GAPDH* gene (ΔC_t_). The ΔΔC_t_ method was used to determine mRNA levels relative to non-transfected negative control samples. Relevant RT-qPCR primer sequences are: h*GAPDH*: 5’ TGCACCACCAACTGCTTAGC 3’ and 5’ ACAGTCTTCTGGGTGGCAGTG 3’. h*CPSF6*: 5’ GATGTGGGTAAAGGAGCAG 3’ and 5’ CTTCATCTGTTGTCCACCA 3’. h*SETD2*: 5’ CTTTCTGTCCCACCCCTGTC 3’ and 5’ CCTTGCACCTCTGATGGCTT 3’. 2-day transfected cells were washed in warm 1X PBS and prepared for circadian synchronization. Synchronization-release medium was pre-warmed and contained 1% v/v FBS, 1.5 μg/ml puromycin, 0.1 mM luciferin, and 0.1 μM dexamethasone. siRNA assays in DMEM were conducted using 2 ml of DMEM (Thermo Fisher # 11966-025) supplemented with 10 mM (low glucose) or 30 mM (high glucose) from a D-glucose stock solution (Sigma # G8644, 100 g/L stock). DMEM base medium contains more total nutrients than MEM—approximately 2-fold higher levels of the 13 essential amino acids, about 4-fold higher levels of the 8 vitamins, and includes the non-essential amino acids (Gly and Ser) in its formulation. siRNA assays in MEM were conducted using 2 ml of MEM (Thermo Fisher # 11095-080) containing 5.56 mM (low glucose) or supplemented up to 25 mM (high glucose) from a D-glucose stock solution (Sigma). Unlike *Neurospora*, complete glucose starvation medium did not support cell viability in preliminary experiments using DMEM medium containing 10% v/v FBS but zero additional glucose. “Low” 5 – 10 mM glucose was defined by manufacturer formulations as well as physiological levels of fasting serum glucose in humans.

### Mammalian luciferase reporter detection and period length calculations

Immediately prior to bioluminescent recording, *Bmal1*-*dLuc* cells in 35-mm dishes were covered with 40-mm circular microscope cover glass (Fisher Scientific # 22038999 40CIR-1) and sealed using high-vacuum silicone grease (Dow Corning # Z273554). Luciferase data were collected in a LumiCycle 32 (ActiMetrics) luminometer every 10 minutes for 5 – 6 days. Raw luciferase traces in bioluminescence counts / second units were exported using LumiCycle analysis software (ActiMetrics, version 2.56). Data from individual plates were manually combined and converted to hours post synchronization using Microsoft Excel. Period lengths for each luciferase trace were calculated using 3 different methods and averaging the period results with equal weights. For the first method, signal peaks and troughs were extracted from days 1 –3 of raw data, and period was estimated by subtracting consecutive peaks or troughs as described (Chen et al., 2020). Second, the WaveClock algorithm was implemented in R (Price et al., 2008). Finally, the suite of *Neurospora* period length tools was used as previously described (Kelliher et al., 2020a). For *Bmal1*-*dLuc* luciferase trace data visualization purposes (Figure 6, Supplementary Figure 6), raw counts per second values sampled within the same hour were averaged together (i.e. data were down-sampled from 10-minute measurement intervals to 1-hour measurement intervals).

### Data visualization

All figures were plotted in R, output as scalable vector graphics, formatted using Inkscape, and archived in R markdown format. Data represent the mean of at least three biological replicates with standard deviation error bars, unless otherwise indicated. All statistical tests were performed in R.

## Supplementary Figures, Tables, and Movies

**Supplementary Figure 1.**
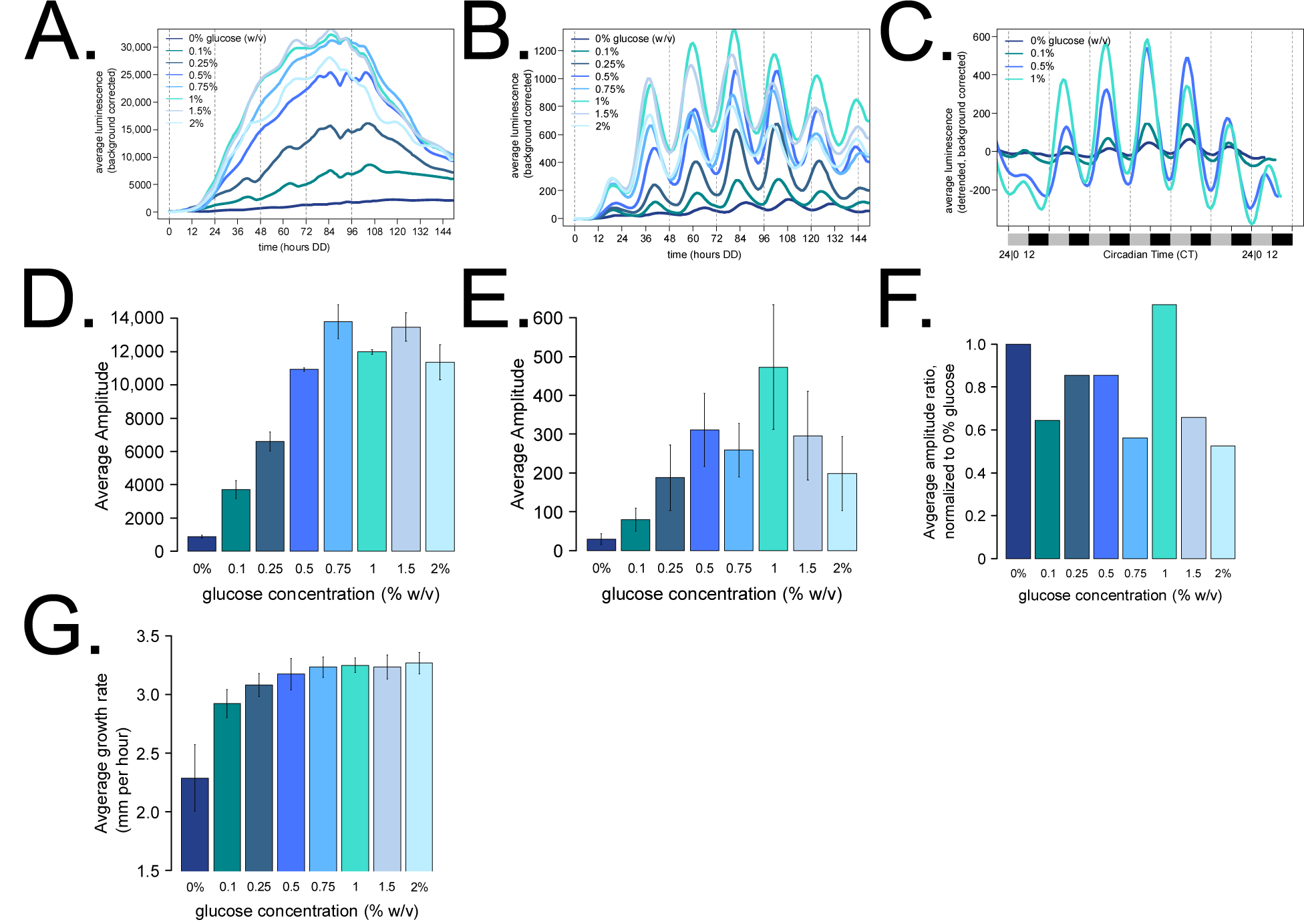
Additional properties of Nutritional Compensation in *Neurospora crassa*. A fungal biomass control was implemented to ask whether the amplitude of the core clock transcriptional reporter changes with glucose levels (e.g. Figure 1B). The *gpd* promoter from *Cochliobolus heterostrophus* driving constitutive *luciferase* was used as a reporter for fungal biomass. Not surprisingly, biomass increases as a function of glucose (N = 3 race tubes per glucose concentration). Many traces showed a decrease in bioluminescence at ∼100 hours, and this correlates with the fungal growth front reaching and surpassing the end of the device’s recording area (**A**) (see: Supplementary Movie 1B). Averaged replicates are shown for the *frq* clock box transcriptional reporter across all glucose levels (N = 6; expanded Figure 1B) (**B**). Detrended clock reporter traces were plotted on a circadian time (CT) scale to normalize for the slight period differences (Figure 1A). Circadian phase is consistent across glucose levels (**C**). Amplitude was computed for each individual biomass reporter trace using data from hours 25 – 108 (amplitude calculation: [maximum value – minimum value] / 2) (**D**). Average amplitude was computed for each individual core clock reporter trace using data from days 2 – 5 (hours 25 – 112) to extract 4 peak and 4 trough values. The first day (hours 0 – 24) was omitted due to low fungal biomass and consequently low luciferase signal during the first recording day (**E**). Core clock amplitudes were normalized to biomass by computing the amplitude ratio at the respective glucose concentrations. There is no clear increasing or decreasing trend of normalized core clock amplitude as a function of glucose, and therefore the higher magnitude oscillations observed at high glucose concentrations are most likely a function of increased biomass only (**F**). Growth rates were computed from biomass reporter experiments by estimating the linear growth rate at 5 consecutive 12-hour intervals from 36 – 84 hours in constant darkness (**G**). No arginine was added to the race tube medium for any experiment shown.

**Supplementary Figure 2.**
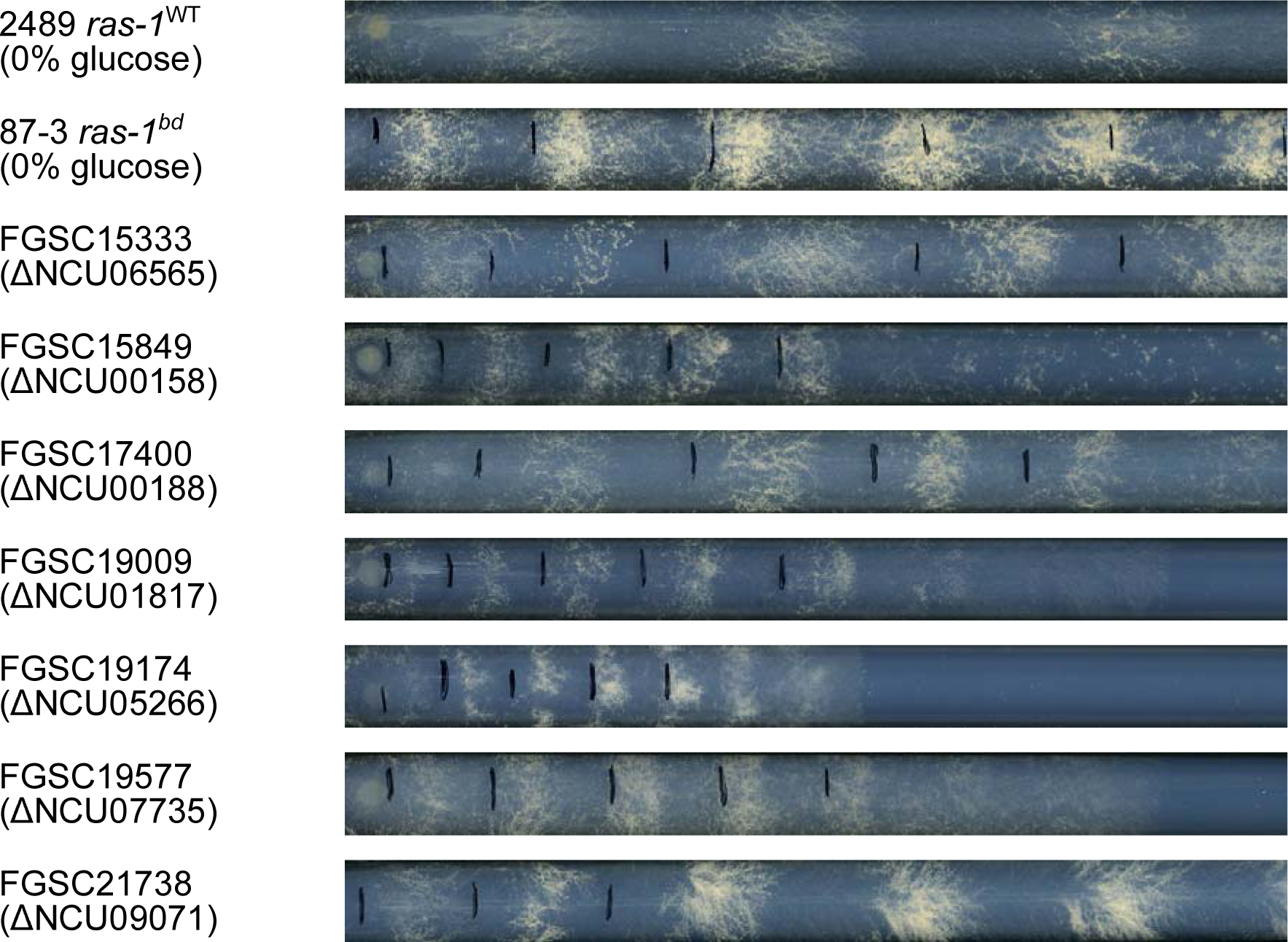
A subset of FGSC knockout strains were identified with strong conidial banding phenotypes on glucose starvation medium. A representative race tube from the primary genetic screen is shown. 6 out of 7 knockout strains with strong banding phenotypes have a wild-type circadian period length and normal compensation, except for FGSC15333 (Figure 3). Like the *ras-1^bd^* (NCU08823) point mutant, all knockout strains have a reduced linear growth rate compared to the wild-type control FGSC2489.

**Supplementary Figure 3.**
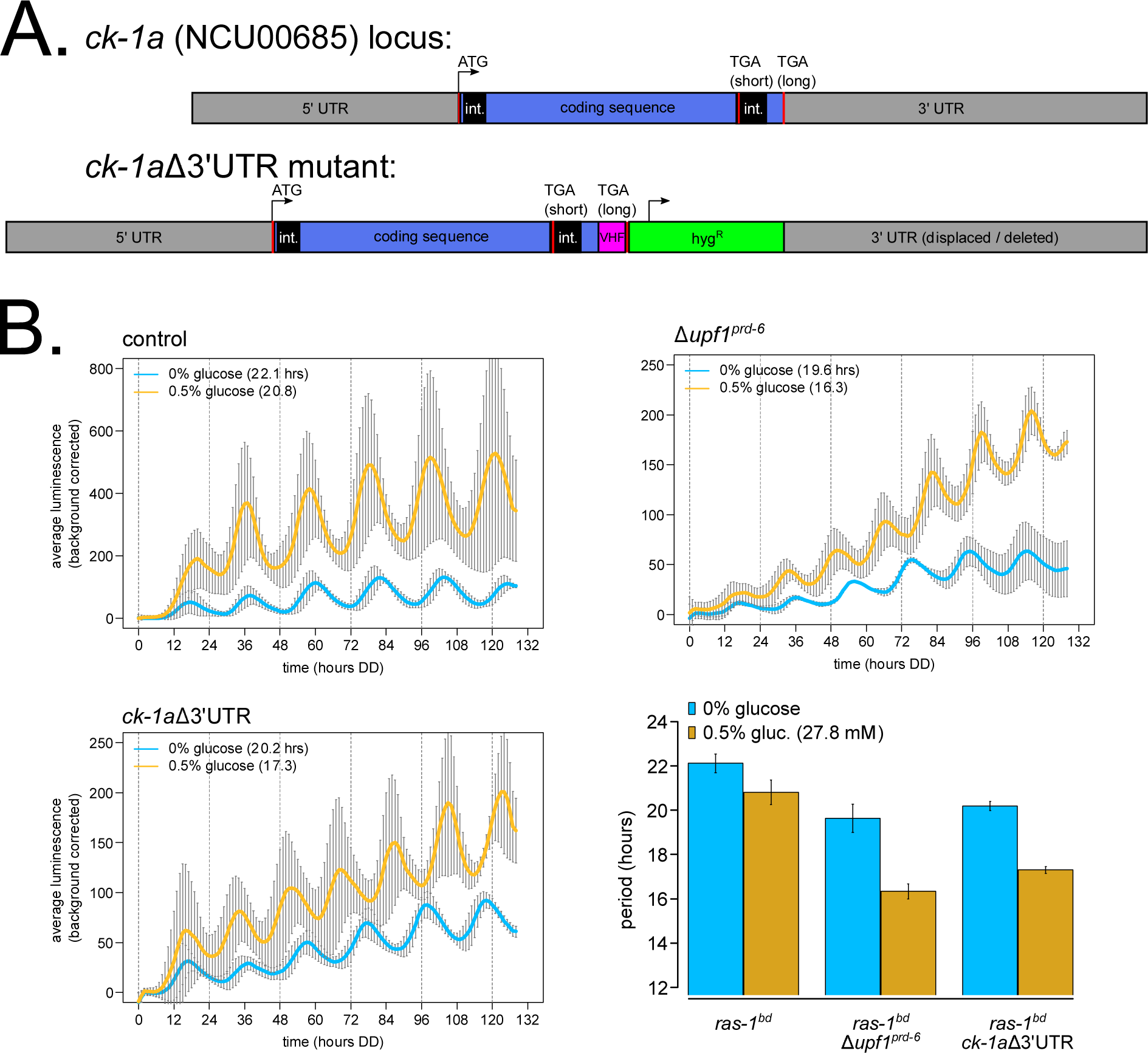
Deletion of the *casein kinase I* 3’ UTR phenocopies the short period length of nonsense-mediated decay (NMD) mutants, and partially explains the nutritional under-compensation phenotype of NMD mutants. The CKI 3’ UTR mutant was constructed by C-terminal epitope-tagging (V5-6xHis-3xFLAG) the LONG isoform of NCU00685 at the endogenous locus and displacing 1,543 bps of the annotated 3’ UTR region. Mutant strain construction is shown with a cartoon diagram to scale (**A**). Circadian bioluminescence was recorded from race tube cultures of the indicated genotypes. High nutrient medium (yellow lines) contained 0.5% w/v glucose 0.17% w/v arginine, and zero nutrient medium (blue lines) contained 0% glucose 0% arginine. Period lengths were computed (N = 3 - 4 biological replicates per nutrient concentration) and summarized in a bar graph (**B**).

**Supplementary Figure 4.**
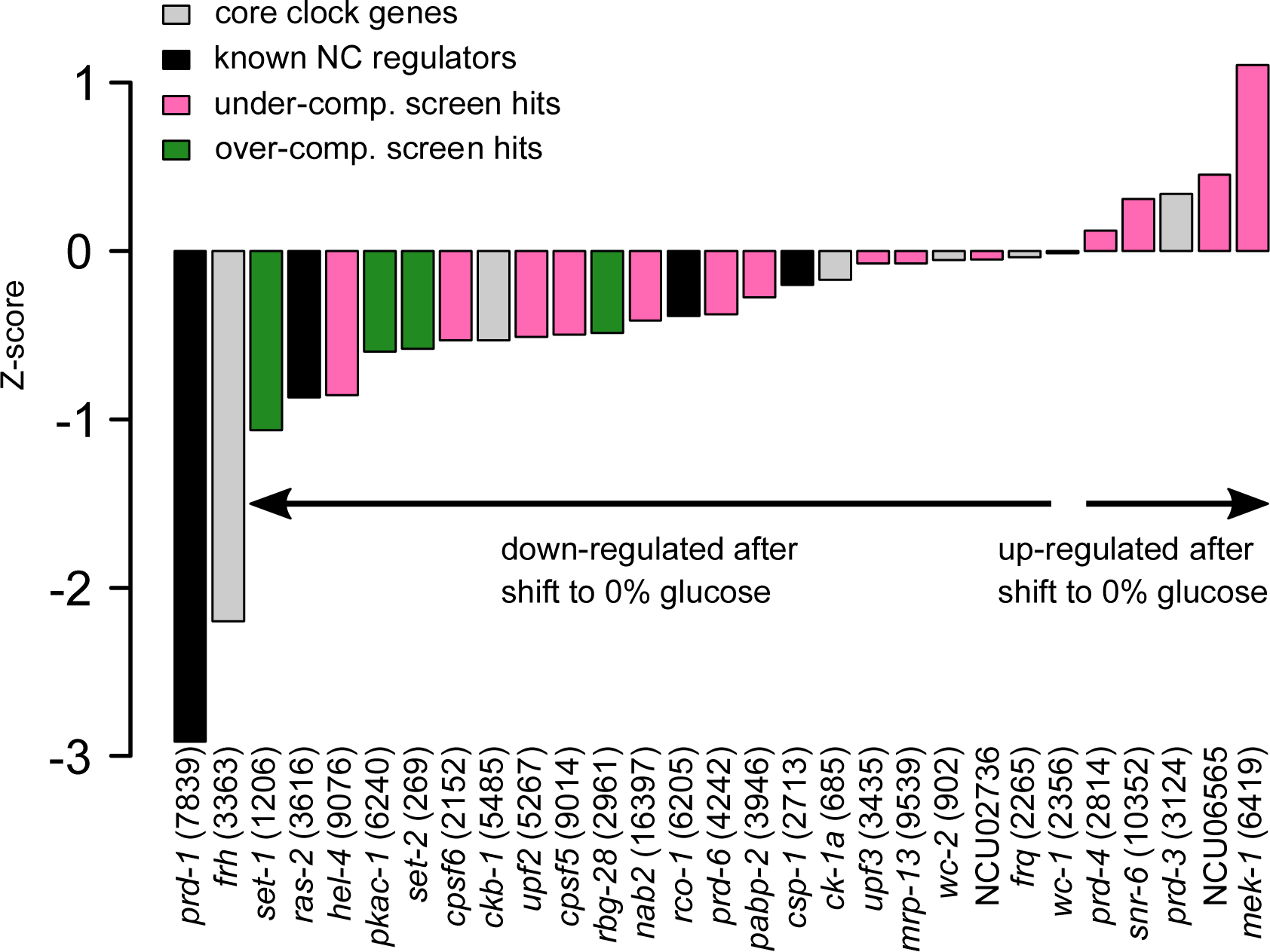
Core clock and compensation gene expression upon glucose starvation. RNA-Sequencing data were mined from a previous study (Wang et al., 2017) (Materials and Methods), where liquid cultures were either maintained in 2% glucose or shifted to glucose starvation for 60 minutes. Duplicate transcriptomes from the 2% glucose condition were compared to 0% glucose starvation replicates, Z-scores were computed for the 8,796 expressed genes in the dataset, and *prd-1* (NCU07839) and *frh* (NCU03363) were found among the top 220 genes (top 2.5%) in the entire transcriptome down-regulated after glucose starvation.

**Supplementary Figure 5.**
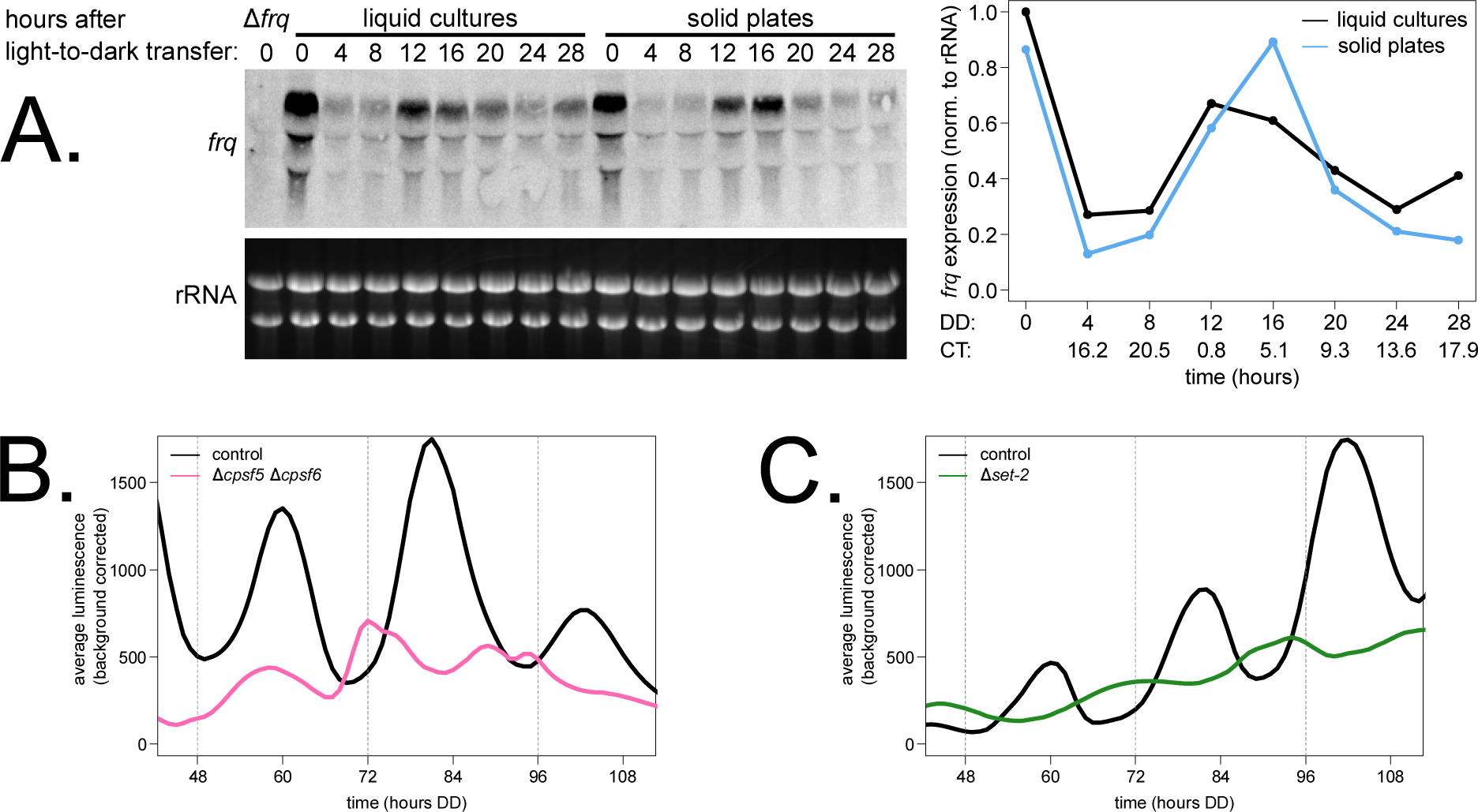
Solid medium cellophane plate cultures maintain circadian function and recapitulate Nutritional Compensation phenotypes of interest. Liquid cultures containing wild-type fungal plugs (1.8% glucose) and cellophane plate cultures inoculated with wild-type conidia (0.5% w/v glucose, 0.17% w/v arginine) were set up concurrently. Liquid and solid cultures were entrained in constant light at 25°C for at least 16 hours, and serial light-to-dark transfers were performed to sample 1.5 cycles of circadian time points from DD4 to DD28 (4-hour sampling density, 44 – 48 hour total culture ages). Total RNA was isolated from each time course sample, and *frq* mRNA rhythms were examined by Northern blot (N = 1 time course replicate). RNA levels were quantified using ImageJ densitometry, normalized, and plotted as line graphs. The circadian clock is clearly functional in both growth regimes (**A**). Circadian bioluminescence was recorded from cellophane plate cultures of the indicated genotypes grown on high nutrient medium (0.25% w/v glucose, 0.17% w/v arginine). One representative luciferase trace is shown from N = 2 biological replicates per strain. Period lengths were calculated, and results agree with Nutritional Compensation phenotypes derived from the race tube screen (Supplementary Table 1): control: 20.6 ± 0.3 hrs; Δ*cpsf5* Δ*cpsf6* double mutant: 17.5 ± 0.4 hrs (**B**). Circadian bioluminescence was recorded from cellophane plate cultures of the indicated genotypes grown on zero nutrient medium (0% glucose, 0% arginine). One representative luciferase trace is shown from N = 2 biological replicates per strain. Period lengths were calculated, and results agree with Nutritional Compensation phenotypes derived from the race tube screen (Supplementary Table 1): control: 22.0 ± 0.6 hrs; Δ*set-2*: 20.5 ± 0.1 hrs (**C**).

**Supplementary Figure 6.**
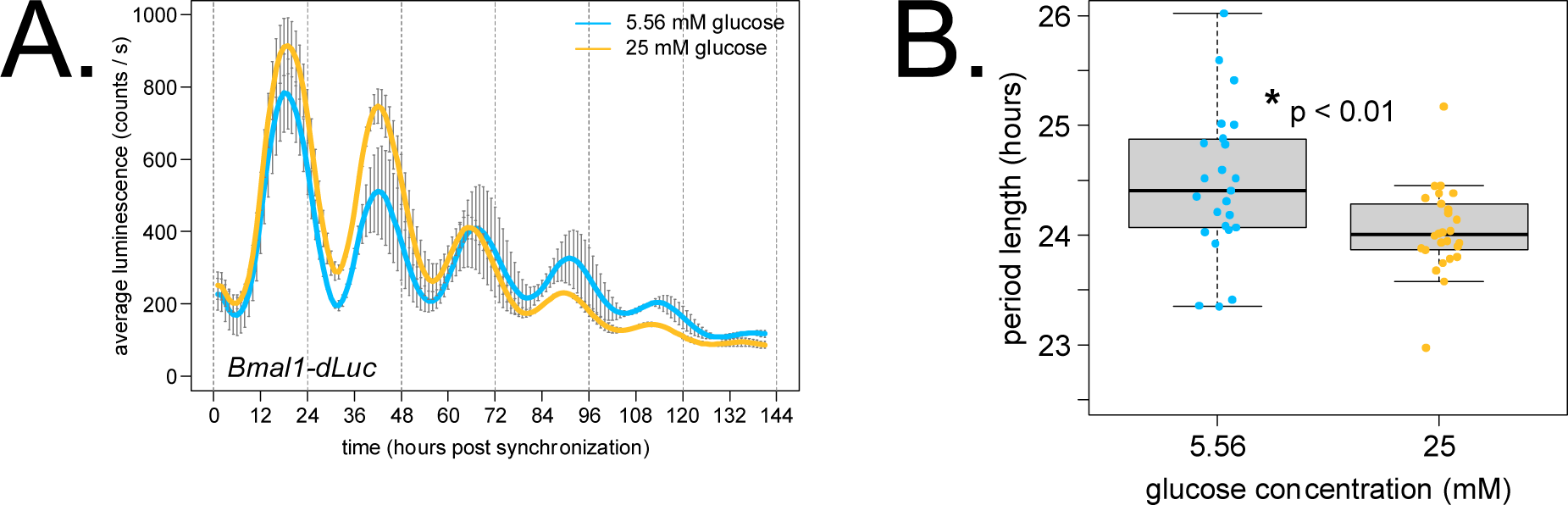
*Bmal1-dLuc* cells are slightly under-compensated from 5.56 mM low glucose to 25 mM high glucose. U2OS cells were synchronized using dexamethasone and released into DMEM low or high glucose medium. Representative luciferase traces are shown from one biological replicate with N = 4 technical replicates per glucose concentration (standard deviation error bars) (**A**). Period lengths were calculated for 25 – 26 total replicates per glucose concentration and plotted as a boxplot. As observed in *Neurospora*, the human circadian period length is under-compensated and shortens slightly with increasing glucose (p < 0.01, student’s t-test). Average period lengths were: 24.5 ± 0.7 hrs (low glucose) and 24.0 ± 0.4 hrs (high glucose) (**B**).

**Supplementary Table 1. Genetic screen for Nutritional Compensation defects**. Results are presented from the 3-phase genetic screen for Nutritional Compensation defects among *Neurospora* knockout strains. Period lengths are shown for each knockout strain. Knockout strains were retained through each phase of the screen if circadian period changes were observed relative to the wild-type control. Nutritional under-compensation mutants were defined by ratios of high-to-zero glucose period lengths ≤ 0.90 (pink font). Nutritional over-compensation mutants had period length ratios ≥ 1.02 (green font).

**Supplementary Table 2. Informatic description of *Neurospora* genes screened for Nutritional Compensation defects**. Circadian rhythmicity at the gene and protein level among knockouts screened was determined from previous studies (Hurley et al., 2014, 2018). Promoter binding by the WCC positive arm transcription factors was determined from previous studies (Smith et al., 2010; Hurley et al., 2014; Sancar et al., 2015). Light-regulated gene activation or repression was determined from previous studies (Chen et al., 2009; Wu et al., 2014; Sancar et al., 2015). Gene expression Z-scores from carbon starvation conditions were also reported (Wang et al., 2017) (Supplementary Figure 4).

**Supplementary Table 3. Consensus list of *Neurospora* genes with Alternative Polyadenylation (APA) in 3’ UTRs**. 843 genes contain multiple poly(A) sites within 3’ UTR regions from the intersection of this study and previous work (Zhou et al., 2018). 9 / 843 genes with 3’ UTR APA are highlighted as core clock genes or compensation screen hits. Annotated MACS2 poly(A) peak results are shown for the 843 genes with Alternative Polyadenylation from each dataset as individual tabs.

**Supplementary Table 4. Alternative Polyadenylation (APA) events altered in the** Δ**CFIm knockout mutant compared to controls**. 1,447 total genes contain multiple poly(A) sites within 3’ UTR regions in wild-type control and/or ΔCFIm mutant data from this study. 940 / 1,447 genes display APA in all 4 datasets. 123 / 1,447 instances are recorded where a single poly(A) peak in control expands to multiple APA events in mutant (dark green highlight). 193 / 1,447 instances are reported of a single poly(A) peak in mutant expanding to multiple APA events in wild-type controls (light green highlight). 155 / 1,447 APA events occur where the location of the predominant poly(A) peak was significantly changed in the mutant background (orange highlight). Annotated MACS2 poly(A) peak results are shown for the APA genes from each dataset as individual tabs.

**Supplementary Movie 1. Nutritional Compensation properties of wild-type *Neurospora crassa*.** Circadian bioluminescence of exemplar wild-type race tubes (Figure 1) is shown for zero nutrient medium (blue line; 0% glucose 0% arginine) (**A**) compared to high nutrient medium (yellow line; 0.5% w/v glucose 0% arginine) (**B**). Average period lengths are 21.9 ± 0.5 hours for 0% glucose and 20.8 ± 0.2 hours for 0.5% w/v glucose (average ± SD).

**Supplementary Movie 2. Nutritional Compensation defect in the** Δ***cpsf6* mutant.** Circadian bioluminescence of one exemplar Δ*cpsf6* race tube (Figure 4A) is shown for whole-tube quantification of high amino acid medium growth (yellow line; 0% glucose 0.17% w/v arginine; average period = 19.4 ± 0.3 hrs) (**A**) compared to old tissue quantification (blue line; 0% glucose 0.17% w/v arginine; period = 17.3 ± 0.3 hrs) (**B**). This ∼2-hour period difference indicates that Δ*cpsf* mutants must undergo a transition from high-to-low amino acid levels to reveal the Nutritional Compensation defect.

## Acknowledgements

This work is supported by the National Institutes of Health (F32 GM128252 to CMK, R35 GM118021 to JCD, and R35 GM118022 to JJL). We thank the Fungal Genetics Stock Center (Kansas City, Missouri, USA) for curating *N. crassa* strains and the whole genome deletion collection (P01 GM068087). We thank Adrienne Mehalow and Arko Dasgupta for generously sharing the collection of *Neurospora* kinase knockout circadian reporter strains, Bradley Bartholomai for the constitutive promoter driving luciferase construct, Consuelo Olivares-Yañez and Alejandra Goity from the Luis Larrondo laboratory at the Pontificia Universidad Católica de Chile for their virtual assistance with the cellophane plate growth method, Lauren Francey from the John Hogenesch laboratory at Cincinnati Children’s Hospital Medical Center for virtual assistance with the U2OS siRNA knockdown experiments, Chris Baker at the Jackson Laboratory for discussions on the *Neurospora* CFIm complex, Josh Gamsby at the University of South Florida for the *Bmal1-dLuc* stable cell line, and Christine Mayr at Memorial Sloan Kettering Cancer Center for discussions and mentorship on the Alternative Polyadenylation (APA) work. 3’ End Sequencing was carried out at the Geisel School of Medicine at Dartmouth in the Genomics Shared Resource (GSR) in collaboration with Fred Kolling IV. Dartmouth GSR is supported by NIH and NSF equipment grants as well as an NCI Cancer Center Core Grant (P30 CA023108).

## References

Abbas M, Moradi F, Hu W, Regudo KL, Osborne M, Pettipas J, Atallah DS, Hachem R, Ott- Peron N, Stuart JA. 2021. Vertebrate cell culture as an experimental approach – limitations and solutions. Comparative Biochemistry and Physiology Part - B: Biochemistry and Molecular Biology, 254. doi:10.1016/j.cbpb.2021.110570

Adhvaryu KK, Morris SA, Strahl BD, Selker EU. 2005. Methylation of histone H3 lysine 36 is required for normal development in *Neurospora crassa*. Eukaryotic Cell, 4(8), 1455–1464. doi:10.1128/EC.4.8.1455-1464.2005

Anders S, Pyl PT, Huber W. 2015. HTSeq-A Python framework to work with high-throughput sequencing data. Bioinformatics, 31(2), 166–169. doi:10.1093/bioinformatics/btu638

Aronson BD, Johnson KA, Loros JJ, Dunlap JC. 1994. Negative feedback defining a circadian clock: autoregulation of the clock gene *frequency*. Science, 263(5153), 1578–84. doi:10.1126/science.8128244

Asher G, Sassone-Corsi P. 2015. Time for food: The intimate interplay between nutrition, metabolism, and the circadian clock. Cell, 161(1), 84–92. doi:10.1016/j.cell.2015.03.015

Baek M, Virgilio S, Lamb TM, Ibarra O, Andrade JM, Gonçalves RD, Dovzhenok A, Lim S, Bell- Pedersen D, Bertolini MC, Hong CI. 2019. Circadian clock regulation of the glycogen synthase (*gsn*) gene by WCC is critical for rhythmic glycogen metabolism in *Neurospora crassa*. Proceedings of the National Academy of Sciences of the United States of America, 116(21), 10435–10440. doi:10.1073/pnas.1815360116

Baggs JE, Price TS, Ditacchio L, Panda S, Fitzgerald GA, Hogenesch JB. 2009. Network features of the mammalian circadian clock. PLOS Biology, 7(3), 0563–0575. doi:10.1371/journal.pbio.1000052

Barrett RK, Takahashi JS. 1995. Temperature compensation and temperature entrainment of the chick pineal cell circadian clock. Journal of Neuroscience, 15(8), 5681–5692. doi:10.1523/jneurosci.15-08-05681.1995

Bartholomai BM. 2021. Cell Biological Dissection of the *Neurospora* Circadian Oscillator: Tools, Approaches, and Fundamental Observations. Ph.D. Thesis, Dartmouth College, 1–207.

Basenko EY, Pulman JA, Shanmugasundram A, Harb OS, Crouch K, Starns D, Warrenfeltz S, Aurrecoechea C, Stoeckert CJ, Kissinger JC, Roos DS, Hertz-Fowler C. 2018. FungiDB: An integrated bioinformatic resource for fungi and oomycetes. Journal of Fungi, 4(1), 1–28. doi:10.3390/jof4010039

Bass J, Takahashi JS. 2010. Circadian integration of metabolism and energetics. Science, 330(6009), 1349–54. doi:10.1126/science.1195027

Beesley S, Kim DW, D’Alessandro M, Jin Y, Lee K, Joo H, Young Y, Tomko RJ, Faulkner J, Gamsby J, Kim JK, Lee C. 2020. Wake-sleep cycles are severely disrupted by diseases affecting cytoplasmic homeostasis. Proceedings of the National Academy of Sciences of the United States of America, 117(45), 28402–28411. doi:10.1073/pnas.2003524117

Belden WJ, Larrondo LF, Froehlich AC, Shi M, Chen CH, Loros JJ, Dunlap JC. 2007. The *band* mutation in *Neurospora crassa* is a dominant allele of *ras-1* implicating RAS signaling in circadian output. Genes & Development, 21(12), 1494–1505. doi:10.1101/gad.1551707

Belden WJ, Loros JJ, Dunlap JC. 2007. Execution of the Circadian Negative Feedback Loop in *Neurospora* Requires the ATP-Dependent Chromatin-Remodeling Enzyme CLOCKSWITCH. Molecular Cell, 25(4), 587–600. doi:10.1016/j.molcel.2007.01.010

Bicocca VT, Ormsby T, Adhvaryu KK, Honda S, Selker EU. 2018. ASH1-catalyzed H3K36 methylation drives gene repression and marks H3K27me2/3-competent chromatin. eLife, 7, 1–19. doi:10.7554/eLife.41497

Cao X, Yang Y, Selby CP, Liu Z, Sancar A. 2021. Molecular mechanism of the repressive phase of the mammalian circadian clock. Proceedings of the National Academy of Sciences of the United States of America, 118(2), 1–9. doi:10.1073/pnas.2021174118

Chen CH, Ringelberg CS, Gross RH, Dunlap JC, Loros JJ. 2009. Genome-wide analysis of light-inducible responses reveals hierarchical light signalling in *Neurospora*. The EMBO Journal, 28(8), 1029–1042. doi:10.1038/emboj.2009.54

Chen S, Fuller KK, Dunlap JC, Loros JJ. 2020. A Pro- and Anti-inflammatory Axis Modulates the Macrophage Circadian Clock. Frontiers in Immunology, 11(May), 9–21. doi:10.3389/fimmu.2020.00867

Cheng P, Yang Y, Liu Y. 2001. Interlocked feedback loops contribute to the robustness of the *Neurospora* circadian clock. Proceedings of the National Academy of Sciences of the United States of America, 98(13), 7408–13. doi:10.1073/pnas.121170298

Collins EJ, Cervantes-Silva MP, Timmons GA, O’Siorain JR, Curtis AM, Hurley JM. 2021. Post-transcriptional circadian regulation in macrophages organizes temporally distinct immunometabolic states. Genome Research, 31(2), 171–185. doi:10.1101/GR.263814.120

Colot HV, Park G, Turner GE, Ringelberg C, Crew CM, Litvinkova L, Weiss RL, Borkovich KA, Dunlap JC. 2006. A high-throughput gene knockout procedure for *Neurospora* reveals functions for multiple transcription factors. Proceedings of the National Academy of Sciences of the United States of America, 103(27), 10352–10357. doi:10.1073/pnas.0601456103

Crosthwaite SK, Dunlap JC, Loros JJ. 1997. *Neurospora wc-1* and *wc-2*: transcription, photoresponses, and the origins of circadian rhythmicity. Science, 276(5313), 763–769. doi:10.1126/science.276.5313.763

Dasgupta A. 2015. Biological Significance of VIVID Photocycle Length and the Control of Gene Expression by the *Neurospora* Clock. Ph.D. Thesis, Dartmouth College, 1–289.

Dibner C, Schibler U. 2015. Circadian timing of metabolism in animal models and humans. Journal of Internal Medicine, 277(5), 513–527. doi:10.1111/joim.12347

Dibner C, Sage D, Unser M, Bauer C, D ’eysmond T, Naef F, Schibler U. 2009. Circadian gene expression is resilient to large fluctuations in overall transcription rates. The EMBO Journal, 28(2), 123–134. doi:10.1038/emboj.2008.262

Dobin A, Davis CA, Schlesinger F, Drenkow J, Zaleski C, Jha S, Batut P, Chaisson M, Gingeras TR. 2013. STAR: Ultrafast universal RNA-seq aligner. Bioinformatics, 29(1), 15–21. doi:10.1093/bioinformatics/bts635

Dovzhenok AA, Baek M, Lim S, Hong CI. 2015. Mathematical modeling and validation of glucose compensation of the *Neurospora* circadian clock. Biophysical Journal, 108(7), 1830–1839. doi:10.1016/j.bpj.2015.01.043

Dunlap JC. 1999. Molecular bases for circadian clocks. Cell, 96(2), 271–290. doi:10.1016/S0092-8674(00)80566-8

Emerson JM, Bartholomai BM, Ringelberg CS, Baker SE, Loros JJ, Dunlap JC. 2015. *period-1* encodes an ATP-dependent RNA helicase that influences nutritional compensation of the *Neurospora* circadian clock. Proceedings of the National Academy of Sciences of the United States of America, 112(51), 15707–15712. doi:10.1073/pnas.1521918112

Etchegaray JP, Yang X, Debruyne JP, Peters AHFM, Weaver DR, Jenuwein T, Reppert SM. 2006. The polycomb group protein EZH2 is required for mammalian circadian clock function. Journal of Biological Chemistry, 281(30), 21209–21215. doi:10.1074/jbc.M603722200

Finger AM, Dibner C, Kramer A. 2020. Coupled network of the circadian clocks: a driving force of rhythmic physiology. FEBS Letters, 594(17), 2734–2769. doi:10.1002/1873-3468.13898

Freitag M. 2017. Histone Methylation by SET Domain Proteins in Fungi. Annual Review of Microbiology, 71, 413–439. doi:10.1146/annurev-micro-102215-095757

Gagliano O, Luni C, Li Y, Angiolillo S, Qin W, Panariello F, Cacchiarelli D, Takahashi JS, Elvassore N. 2021. Synchronization between peripheral circadian clock and feeding-fasting cycles in microfluidic device sustains oscillatory pattern of transcriptome. Nature Communications, 12(1), 1–12. doi:10.1038/s41467-021-26294-9

Gamsby JJ, Loros JJ, Dunlap JC. 2009. A phylogenetically conserved DNA damage response resets the circadian clock. Journal of Biological Rhythms, 24(3), 193–202. doi:10.1177/0748730409334748

Gardner GF, Feldman JF. 1981. Temperature Compensation of Circadian Period Length in Clock Mutants of *Neurospora crassa*. Plant Physiology, 68(6), 1244–8.

Gendreau KL, Unruh BA, Zhou C, Kojima S. 2018. Identification and characterization of transcripts regulated by circadian alternative polyadenylation in mouse liver. G3: Genes, Genomes, Genetics, 8(11), 3539–3548. doi:10.1534/g3.118.200559

Greenwell BJ, Beytebiere JR, Lamb TM, Bell-Pedersen D, Merlin C, Menet JS. 2020. Isoform-specific regulation of rhythmic gene expression by alternative polyadenylation. bioRxiv, 1, 1–28. doi:10.1101/2020.12.12.422514

Gyöngyösi N, Szöke A, Ella K, Káldi K. 2017. The small G protein RAS2 is involved in the metabolic compensation of the circadian clock in the circadian model *Neurospora crassa*. Journal of Biological Chemistry, 292(36), 14929–14939. doi:10.1074/jbc.M117.804922

Hastings JW, Sweeney BM. 1957. On the Mechanism of Temperature Independence in a Biological Clock. Proceedings of the National Academy of Sciences of the United States of America, 43(9), 804–811. doi:10.1073/pnas.43.9.804

Hatori M, Vollmers C, Zarrinpar A, DiTacchio L, Bushong EA, Gill S, Leblanc M, Chaix A, Joens M, Fitzpatrick JAJ, Ellisman MH, Panda S. 2012. Time-restricted feeding without reducing caloric intake prevents metabolic diseases in mice fed a high-fat diet. Cell Metabolism, 15(6), 848–860. doi:10.1016/j.cmet.2012.04.019

Hogan GJ, Brown PO, Herschlag D. 2015. Evolutionary Conservation and Diversification of Puf RNA Binding Proteins and Their mRNA Targets. PLOS Biology, 13(11), 1–47. doi:10.1371/journal.pbio.1002307

Hong L, Lavrentovich DO, Chavan A, Leypunskiy E, Li E, Matthews C, LiWang A, Rust MJ, Dinner AR. 2020. Bayesian modeling reveals metabolite-dependent ultrasensitivity in the cyanobacterial circadian clock. Molecular Systems Biology, 16(6), 1–23. doi:10.15252/msb.20199355

Hu Y, Liu X, Lu Q, Yang Y, He Q, Liu Y, Liu X. 2021. FRQ-CK1 interaction underlies temperature compensation of the *Neurospora* circadian clock. mBio, 12(3). doi:10.1128/mBio.01425-21

Huberman LB, Wu VW, Kowbel DJ, Lee J, Daum C, Grigoriev I V., O’Malley RC, Louise Glass N. 2021. DNA affinity purification sequencing and transcriptional profiling reveal new aspects of nitrogen regulation in a filamentous fungus. Proceedings of the National Academy of Sciences of the United States of America, 118(13), 1–11. doi:10.1073/pnas.2009501118

Hurley JM, Dasgupta A, Emerson JM, Zhou X, Ringelberg CS, Knabe N, Lipzen AM, Lindquist EA, Daum CG, Barry KW, Grigoriev I V., Smith KM, Galagan JE, Bell-Pedersen D, Freitag M, Cheng C, Loros JJ, Dunlap JC. 2014. Analysis of clock-regulated genes in *Neurospora* reveals widespread posttranscriptional control of metabolic potential. Proceedings of the National Academy of Sciences of the United States of America, 111(48), 16995–17002. doi:10.1073/pnas.1418963111

Hurley JM, Jankowski MS, De Los Santos H, Crowell AM, Fordyce SB, Zucker JD, Kumar N, Purvine SO, Robinson EW, Shukla A, Zink E, Cannon WR, Baker SE, Loros JJ, Dunlap JC. 2018. Circadian Proteomic Analysis Uncovers Mechanisms of Post-Transcriptional Regulation in Metabolic Pathways. Cell Systems, 7(6), 613–626.e5. doi:10.1016/j.cels.2018.10.014

Hurley JM, Larrondo LF, Loros JJ, Dunlap JC. 2013. Conserved RNA helicase FRH acts nonenzymatically to support the intrinsically disordered *Neurospora* clock protein FRQ. Molecular Cell, 52(6), 832–843. doi:10.1016/j.molcel.2013.11.005

Izumo M, Johnson CH, Yamazaki S. 2003. Circadian gene expression in mammalian fibroblasts revealed by real-time luminescence reporting: temperature compensation and damping. Proceedings of the National Academy of Sciences of the United States of America, 100(26), 16089–94. doi:10.1073/pnas.2536313100

Johnson CH, Egli M. 2014. Metabolic compensation and circadian resilience in prokaryotic cyanobacteria. Annual Review of Biochemistry, 83, 221–247. doi:10.1146/annurev-biochem-060713-035632

Johnson CH, Elliott JA, Foster R. 2003. Entrainment of circadian programs. Chronobiology International, 20(5), 741–774. doi:10.1081/CBI-120024211

Ju D, Zhang W, Yan J, Zhao H, Li W, Wang J, et al. 2020. Chemical perturbations reveal that RUVBL2 regulates the circadian phase in mammals. Science Translational Medicine, 12(542), 1–12. doi:10.1126/scitranslmed.aba0769

Karki S, Castillo K, Ding Z, Kerr O, Lamb TM, Wu C, Sachs MS, Bell-Pedersen D. 2020. Circadian clock control of eIF2α phosphorylation is necessary for rhythmic translation initiation. Proceedings of the National Academy of Sciences of the United States of America, 117(20), 10935–10945. doi:10.1073/pnas.1918459117

Katada S, Sassone-Corsi P. 2010. The histone methyltransferase MLL1 permits the oscillation of circadian gene expression. Nature Structural and Molecular Biology, 17(12), 1414–1421. doi:10.1038/nsmb.1961

Kelliher CM, Lambreghts R, Xiang Q, Baker CL, Loros JJ, Dunlap JC. 2020. Prd-2 directly regulates *casein kinase I* and counteracts nonsense-mediated decay in the *Neurospora* circadian clock. eLife, 9, 1–22. doi:10.7554/ELIFE.64007

Kelliher CM, Loros JJ, Dunlap JC. 2020. Evaluating the circadian rhythm and response to glucose addition in dispersed growth cultures of *Neurospora crassa*. Fungal Biology, 124(5), 398–406. doi:10.1016/j.funbio.2019.11.004

Kidd PB, Young MW, Siggia ED. 2015. Temperature compensation and temperature sensation in the circadian clock. Proceedings of the National Academy of Sciences of the United States of America, 112(46), E6284–E6292. doi:10.1073/pnas.1511215112

Kohsaka A, Laposky AD, Ramsey KM, Estrada C, Joshu C, Kobayashi Y, Turek FW, Bass J. 2007. High-Fat Diet Disrupts Behavioral and Molecular Circadian Rhythms in Mice. Cell Metabolism, 6(5), 414–421. doi:10.1016/j.cmet.2007.09.006

Koike N, Yoo S-H, Huang H-C, Kumar V, Lee C, Kim T-K, Takahashi JS. 2012. Transcriptional architecture and chromatin landscape of the core circadian clock in mammals. Science, 338(6105), 349–54. doi:10.1126/science.1226339

Krishnaiah SY, Wu G, Altman BJ, Growe J, Rhoades SD, Coldren F, et al. 2017. Clock Regulation of Metabolites Reveals Coupling between Transcription and Metabolism. Cell Metabolism, 25(5), 1206. doi:10.1016/j.cmet.2017.04.023

Kubo T, Wada T, Yamaguchi Y, Shimizu A, Handa H. 2006. Knock-down of 25 kDa subunit of cleavage factor Im in Hela cells alters alternative polyadenylation within 3 -UTRs. Nucleic Acids Research, 34(21), 6264–6271. doi:10.1093/nar/gkl794

Lambreghts R, Shi M, Belden WJ, DeCaprio D, Park D, Henn MR, Galagan JE, Batürkmen M, Birren BW, Sachs MS, Dunlap JC, Loros JJ. 2009. A high-density single nucleotide polymorphism map for *Neurospora crassa*. Genetics, 181(2), 767–781. doi:10.1534/genetics.108.089292

Larrondo LF, Loros JJ, Dunlap JC. 2012. High-resolution spatiotemporal analysis of gene expression in real time: In vivo analysis of circadian rhythms in *Neurospora crassa* using a FREQUENCY-luciferase translational reporter. Fungal Genetics and Biology, 49(9), 681–683. doi:10.1016/j.fgb.2012.06.001

Larrondo LF, Olivares-Yañez C, Baker CL, Loros JJ, Dunlap JC. 2015. Decoupling circadian clock protein turnover from circadian period determination. Science, 347(6221). doi:10.1126/science.1257277

Lee Y, Shen Y, Francey LJ, Ramanathan C, Sehgal A, Liu AC, Hogenesch JB. 2019. The NRON complex controls circadian clock function through regulated PER and CRY nuclear translocation. Scientific Reports, 9(1), 1–12. doi:10.1038/s41598-019-48341-8

Liu X, Blaženović I, Contreras AJ, Pham TM, Tabuloc CA, Li YH, Ji J, Fiehn O, Chiu JC. 2021. Hexosamine biosynthetic pathway and O-GlcNAc-processing enzymes regulate daily rhythms in protein O-GlcNAcylation. Nature Communications, 12(1), 1–16. doi:10.1038/s41467-021-24301-7

Liu X, Li H, Liu Q, Niu Y, Hu Q, Deng H, Cha J, Wang Y, Liu Y, He Q. 2015. Role for Protein Kinase A in the *Neurospora* Circadian Clock by Regulating White Collar-Independent *frequency* Transcription through Phosphorylation of RCM-1. Molecular and Cellular Biology, 35(12), 2088–102. doi:10.1128/MCB.00709-14

Liu Y, Hu W, Murakawa Y, Yin J, Wang G, Landthaler M, Yan J. 2013. Cold-induced RNA-binding proteins regulate circadian gene expression by controlling alternative polyadenylation. Scientific Reports, 3, 1–11. doi:10.1038/srep02054

Loros JJ. 2020. Principles of the animal molecular clock learned from *Neurospora*. European Journal of Neuroscience, 51(1), 19–33.

Lowrey PL, Shimomura K, Antoch MP, Yamazaki S, Zemenides PD, Ralph MR, Menaker M, Takahashi JS. 2000. Positional syntenic cloning and functional characterization of the mammalian circadian mutation tau. Science, 288(5465), 483–92. doi:10.1126/science.288.5465.483

Maier B, Wendt S, Vanselow JT, Wallaeh T, Reischl S, Oehmke S, Sehlosser A, Kramer A. 2009. A large-scale functional RNAi screen reveals a role for CK2 in the mammalian circadian clock. Genes and Development, 23(6), 708–718. doi:10.1101/gad.512209

Marcheva B, Ramsey KM, Buhr ED, Kobayashi Y, Su H, Ko CH, Ivanova G, Omura C, Mo S, Vitaterna MH, Lopez JP, Philipson LH, Bradfield CA, Crosby SD, JeBailey L, Wang X, Takahashi JS, Bass J. 2010. Disruption of the clock components CLOCK and BMAL1 leads to hypoinsulinaemia and diabetes. Nature, 466(7306), 627–31. doi:10.1038/nature09253

Martin G, Gruber AR, Keller W, Zavolan M. 2012. Genome-wide Analysis of Pre-mRNA 3’ End Processing Reveals a Decisive Role of Human Cleavage Factor I in the Regulation of 3’ UTR Length. Cell Reports, 1(6), 753–763. doi:10.1016/j.celrep.2012.05.003

Matsumura R, Okamoto A, Node K, Akashi M. 2014. Compensation for intracellular environment in expression levels of mammalian circadian clock genes. Scientific Reports, 4, 1–6. doi:10.1038/srep04032

Mayr C. 2017. Regulation by 3 -Untranslated Regions. Annual Review of Genetics, 51, 171–′194. doi:10.1146/annurev-genet-120116-024704

Mayr C, Bartel DP. 2009. Widespread Shortening of 3′ Polyadenylation Activates Oncogenes in Cancer Cells. Cell, 138(4), 673–684. doi:10.1016/j.cell.2009.06.016

Mehra A, Shi M, Baker CL, Colot HV, Loros JJ, Dunlap JC. 2009. A Role for Casein Kinase 2 in the Mechanism Underlying Circadian Temperature Compensation. Cell, 137(4), 749–760. doi:10.1016/j.cell.2009.03.019

Menet JS, Pescatore S, Rosbash M. 2014. BMAL1 is a pioneer- like transcription factor. Genes & Development, 28, 8–13. doi:10.1101/gad.228536.113.mouse

Moqtaderi Z, Geisberg J V., Struhl K. 2022. A compensatory link between cleavage/polyadenylation and mRNA turnover regulates steady-state mRNA levels in yeast. Proceedings of the National Academy of Sciences of the United States of America, 119(4), 1–6. doi:10.1073/pnas.2121488119

Muñoz-Guzmán F, .Caballero .V., L.F. Larrondo 2021. A global search for novel transcription factors impacting the *Neurospora crassa* circadian clock. G3: Genes, Genomes, Genetics, 11(6). doi:10.1093/g3journal/jkab100

Nakashima H. 1981. A liquid culture method for the biochemical analysis of the circadian clock of *Neurospora crassa*. Plant & Cell Physiology, 22(2), 231–238.

Olivares-Yanez C, Emerson J, Kettenbach A, Loros JJ, Dunlap JC, Larrondo LF. 2016. Modulation of circadian gene expression and metabolic compensation by the RCO-1 corepressor of *Neurospora crassa*. Genetics, 204(1), 163–176. doi:10.1534/genetics.116.191064

Padmanabhan K, Robles MS, Westerling T, Weitz CJ. 2012. Feedback Regulation of Transcriptional Termination by the Mammalian Circadian Clock PERIOD Complex. Science, 337(6094), 599–602. doi:10.1126/science.1221592

Philpott JM, Narasimamurthy R, Ricci CG, Freeberg AM, Hunt SR, Yee LE, Pelofsky RS, Tripathi S, Virshup DM, Partch CL. 2020. Casein kinase 1 dynamics underlie substrate selectivity and the PER2 circadian phosphoswitch. eLife, 9, 1–28. doi:10.7554/eLife.52343

Philpott JM, Torgrimson MR, Harold RL, Partch CL. 2021. Biochemical mechanisms of period control within the mammalian circadian clock. Seminars in Cell and Developmental Biology. doi:10.1016/j.semcdb.2021.04.012

Phong C, Markson JS, Wilhoite CM, Rust MJ. 2013. Robust and tunable circadian rhythms from differentially sensitive catalytic domains. Proceedings of the National Academy of Sciences of the United States of America, 110(3), 1124–1129. doi:10.1073/pnas.1212113110

Pittendrigh CS, Bruce VG, Rosenweig NS, Rubin ML. 1959. A Biological Clock in *Neurospora*. Nature, 184(4681), 169–170. doi:10.1038/184169a0

Pittendrigh CS, Caldarola PC. 1973. General homeostasis of the frequency of circadian oscillations. Proceedings of the National Academy of Sciences of the United States of America, 70(9), 2697–2701. doi:10.1073/pnas.70.9.2697

Portolés S, Más P. 2010. The functional interplay between protein kinase CK2 and cca1 transcriptional activity is essential for clock temperature compensation in *Arabidopsis*. PLoS Genetics, 6(11). doi:10.1371/journal.pgen.1001201

Price TS, Baggs JE, Curtis AM, Fitzgerald GA, Hogenesch JB. 2008. WAVECLOCK: Wavelet analysis of circadian oscillation. Bioinformatics, 24(23), 2794–2795. doi:10.1093/bioinformatics/btn521

Raduwan H, Isola AL, Belden WJ. 2013. Methylation of histone H3 on lysine 4 by the lysine methyltransferase SET1 protein is needed for normal clock gene expression. Journal of Biological Chemistry, 288(12), 8380–8390. doi:10.1074/jbc.M112.359935

Radziuk J, Pye S. 2006. Diurnal rhythm in endogenous glucose production is a major contributor to fasting hyperglycaemia in type 2 diabetes. Suprachiasmatic deficit or limit cycle behaviour? Diabetologia, 49(7), 1619–1628. doi:10.1007/s00125-006-0273-9

Ralph MR, Menaker M. 1988. A mutation of the circadian system in golden hamsters. Science, 241(4870), 1225–1227. doi:10.1126/science.3413487

Ramanathan C, Kathale ND, Liu D, Lee C, Freeman DA, Hogenesch JB, Cao R, Liu AC. 2018. mTOR signaling regulates central and peripheral circadian clock function. PLoS Genetics, 14(5), 1–21. doi:10.1371/journal.pgen.1007369

Ramanathan C, Xu H, Khan SK, Shen Y, Gitis PJ, Welsh DK, Hogenesch JB, Liu AC. 2014. Cell Type-Specific Functions of Period Genes Revealed by Novel Adipocyte and Hepatocyte Circadian Clock Models. PLoS Genetics, 10(4). doi:10.1371/journal.pgen.1004244

Ray D, Kazan H, Cook KB, Weirauch MT, Najafabadi HS, Li X, et al. 2013. A compendium of RNA-binding motifs for decoding gene regulation. Nature, 499(7457), 172–177. doi:10.1038/nature12311

Roenneberg T, Merrow M. 1999. Circadian systems and metabolism. Journal of Biological Rhythms, 14(6), 449–459. doi:10.1177/074873099129001019

Ruoff P, Behzadi A, Hauglid M, Vinsjevik M, Havas H. 2000. pH homeostasis of the circadian sporulation rhythm in clock mutants of *Neurospora crassa*. Chronobiology International, 17(6), 733–750. doi:10.1081/CBI-100102109

Sancar C, Ha N, Yilmaz R, Tesorero R, Fisher T, Brunner M, Sancar G. 2015. Combinatorial Control of Light Induced Chromatin Remodeling and Gene Activation in *Neurospora*. PLOS Genetics, 11(3), 1–26. doi:10.1371/journal.pgen.1005105

Sancar G, Brunner M. 2014. Circadian clocks and energy metabolism. Cellular and Molecular Life Sciences, 71(14), 2667–2680. doi:10.1007/s00018-014-1574-7

Sancar G, Sancar C, Brunner M. 2012. Metabolic compensation of the *Neurospora* clock by a glucose-dependent feedback of the circadian repressor CSP1 on the core oscillator. Genes & Development, 26(21), 2435–2442. doi:10.1101/gad.199547.112

Sargent ML, Briggs W, Woodward D. 1966. Circadian nature of a rhythm expressed by an invertaseless strain of *Neurospora crassa*. Plant physiology, 41(8), 1343–1349. doi:10.1104/pp.41.8.1343

Sargent ML, Kaltenborn SH. 1972. Effects of Medium Composition and Carbon Dioxide on Circadian Conidiation in *Neurospora*. Plant Physiology, 50(1), 171–175. doi:10.1104/pp.50.1.171

Sato TK, Yamada RG, Ukai H, Baggs JE, Miraglia LJ, Kobayashi TJ, Welsh DK, Kay SA, Ueda HR, Hogenesch JB. 2006. Feedback repression is required for mammalian circadian clock function. Nature Genetics, 38(3), 312–319. doi:10.1038/ng1745

Schmal C, Maier B, Ashwal-Fluss R, Bartok O, Finger A-M, Bange T, Koutsouli S, Robles MS, Kadener S, Herzel H, Kramer A. 2021. An integrative omics approach reveals posttranscriptional mechanisms underlying circadian temperature compensation. bioRxiv, 2021.10.06.463236.

Shapiro ET, Polonsky KS, Copinschi G, Bosson D, Tillil H, Blackman J, Lewis G, Van Cauter E. 1991. Nocturnal elevation of glucose levels during fasting in noninsulin-dependent diabetes. Journal of Clinical Endocrinology and Metabolism, 72(2), 444–454. doi:10.1210/jcem-72-2-444

Shinohara Y, Koyama YM, Ukai-Tadenuma M, Hirokawa T, Kikuchi M, Yamada RG, Ukai H, Fujishima H, Umehara T, Tainaka K, Ueda HR. 2017. Temperature-Sensitive Substrate and Product Binding Underlie Temperature-Compensated Phosphorylation in the Clock. Molecular Cell, 67(5), 783–798.e20. doi:10.1016/j.molcel.2017.08.009

Sinturel F, Gerber A, Mauvoisin D, Wang J, Gatfield D, Stubblefield JJ, Green CB, Gachon F, Schibler U. 2017. Diurnal Oscillations in Liver Mass and Cell Size Accompany Ribosome Assembly Cycles. Cell, 169(4), 651–663.e14. doi:10.1016/j.cell.2017.04.015

Smith KM, Sancar G, Dekhang R, Sullivan CM, Li S, Tag AG, Sancar C, Bredeweg EL, Priest HD, McCormick RF, Thomas TL, Carrington JC, Stajich JE, Bell-Pedersen D, Brunner M, Freitag M. 2010. Transcription factors in light and circadian clock signaling networks revealed by genomewide mapping of direct targets for *Neurospora* white collar complex. Eukaryotic Cell, 9(10), 1549–1556. doi:10.1128/EC.00154-10

Sun G, Zhou Z, Liu X, Gai K, Liu Q, Cha J, Kaleri FN, Wang Y, He Q. 2016. Suppression of WHITE COLLAR-independent *frequency* transcription by histone H3 lysine 36 methyltransferase SET-2 is necessary for clock function in *Neurospora*. Journal of Biological Chemistry, 291(21), 11055–11063. doi:10.1074/jbc.M115.711333

Takahashi JS. 2017. Transcriptional architecture of the mammalian circadian clock. Nature Reviews Genetics, 18(3), 164–179. doi:10.1038/nrg.2016.150

Thurley K, Herbst C, Wesener F, Koller B, Wallach T, Maier B, Kramer A, Westermark PO. 2017. Principles for circadian orchestration of metabolic pathways. Proceedings of the National Academy of Sciences, 114(7), 1572–1577. doi:10.1073/pnas.1613103114

Tosini G, Menaker M. 1998. The tau mutation affects temperature compensation of hamster retinal circadian oscillators. Neuroreport, 9(6), 1001–1005.

Tsuchiya Y, Akashi M, Nishida E. 2003. Temperature compensation and temperature resetting of circadian rhythms in mammalian cultured fibroblasts. Genes to Cells, 8(8), 713–720. doi:10.1046/j.1365-2443.2003.00669.x

Ueda HR, Chen W, Adachi A, Wakamatsu H, Hayashi S, Takasugi T, Nagano M, Nakahama KI, Suzuki Y, Sugano S, Lino M, Shigeyoshi Y, Hashimoto S. 2002. A transcription factor response element for gene expression during circadian night. Nature, 418(6897), 534–539. doi:10.1038/nature00906

Upadhyay A, Brunner M, Herzel H. 2019. An inactivation switch enables rhythms in a *Neurospora* clock model. International Journal of Molecular Sciences, 20(12). doi:10.3390/ijms20122985

Valekunja UK, Edgar RS, Oklejewicz M, Van Der Horst GTJ, O’Neill JS, Tamanini F, Turner DJ, Reddy AB. 2013. Histone methyltransferase MLL3 contributes to genome-scale circadian transcription. Proceedings of the National Academy of Sciences of the United States of America, 110(4), 1554–1559. doi:10.1073/pnas.1214168110

Van Hoof A, Lennertz P, Parker R. 2000. Three conserved members of the RNase D family have unique and overlapping functions in the processing of 5S, 5.8S, U4, U5. RNase MRP and RNase P RNAs in yeast. EMBO Journal, 19(6), 1357–1365. doi:10.1093/emboj/19.6.1357

Wang B, Li J, Gao J, Cai P, Han X, Tian C. 2017. Identification and characterization of the glucose dual-affinity transport system in *Neurospora crassa*: pleiotropic roles in nutrient transport, signaling, and carbon catabolite repression. Biotechnology for Biofuels, 10(1), 17. doi:10.1186/s13068-017-0705-4

Wang B, Kettenbach AN, Gerber SA, Loros JJ, Dunlap JC. 2014. *Neurospora* WC-1 Recruits SWI/SNF to Remodel *frequency* and Initiate a Circadian Cycle. PLOS Genetics, 10(9), e1004599. doi:10.1371/journal.pgen.1004599

Wang B, Kettenbach AN, Zhou X, Loros JJ, Dunlap JC. 2019. The Phospho-Code Determining Circadian Feedback Loop Closure and Output in *Neurospora*. Molecular Cell, 74(4), 771–784. doi:10.1016/j.molcel.2019.03.003

Wehrens SMT, Christou S, Isherwood C, Middleton B, Gibbs MA, Archer SN, Skene DJ, Johnston JD. 2017. Meal Timing Regulates the Human Circadian System. Current Biology, 27(12), 1768–1775.e3. doi:10.1016/j.cub.2017.04.059

Wu C, Yang F, Smith KM, Peterson M, Dekhang R, Zhang Y, Zucker J, Bredeweg EL, Mallappa C, Zhou X, Lyubetskaya A, Townsend JP, Galagan JE, Freitag M, Dunlap JC, Bell- Pedersen D, Sachs MS. 2014. Genome-Wide Characterization of Light-Regulated Genes in *Neurospora crassa*. G3: Genes|Genomes|Genetics, 4(9), 1731–1745. doi:10.1534/g3.114.012617

Wu Y, Zhang Y, Sun Y, Yu J, Wang P, Chen S, Zhang D, He Q, Guo J. 2017. Up-Frameshift Protein UPF1 Regulates *Neurospora crassa* Circadian and Diurnal Growth Rhythms. Genetics, 206, 1881–1893.

Yang Y, Li Y, Sancar A, Oztas O. 2020. The circadian clock shapes the *Arabidopsis* transcriptome by regulating alternative splicing and alternative polyadenylation. The Journal of Biological Chemistry. doi:10.1074/jbc.RA120.013513

Young ATL, Moore RB, Murray AG, Mullen JC, Lakey JRT. 2004. Assessment of Different Transfection Parameters in Efficiency Optimization. Cell Transplantation, 13(2), 179–185. doi:10.3727/000000004773301861

Zaveri K, Kiranmayi P, Nagamruta M, Rachel KV. 2017. RNA-binding Proteins in *Neurospora crassa*: An Insight into Spliceosome and Post Transcriptional Gene Silencing Machinery. Current Proteomics, 14(4), 261–276. doi:10.2174/1570164614666170329145814

Zhang EE, Liu AC, Hirota T, Miraglia LJ, Welch G, Pongsawakul PY, Liu X, Atwood A, Huss JW, Janes J, Su AI, Hogenesch JB, Kay SA. 2009. A Genome-wide RNAi Screen for Modifiers of the Circadian Clock in Human Cells. Cell, 139(1), 199–210. doi:10.1016/j.cell.2009.08.031

Zhang R, Lahens NF, Ballance HI, Hughes ME, Hogenesch JB. 2014. A circadian gene expression atlas in mammals: implications for biology and medicine. Proceedings of the National Academy of Sciences of the United States of America, 111(45), 16219–24. doi:10.1073/pnas.1408886111

Zhang Y, Iiams SE, Menet JS, Hardin PE, Merlin C. 2022. TRITHORAX-dependent arginine methylation of HSP68 mediates circadian repression by PERIOD in the monarch butterfly. Proceedings of the National Academy of Sciences of the United States of America, 119(4), 1–10. doi:10.1073/pnas.2115711119

Zhang Y, Liu T, Meyer CA, Eeckhoute J, Johnson DS, Bernstein BE, Nussbaum C, Myers RM, Brown M, Li W, Liu XS. 2008. Model-based Analysis of ChIP-Seq (MACS). Genome Biology, 9(9), R137. doi:10.1186/gb-2008-9-9-r137

Zhou Z, Dang Y, Zhou M, Yuan H, Liu Y. 2018. Codon usage biases co-evolve with transcription termination machinery to suppress premature cleavage and polyadenylation. eLife, 7, 1–29. doi:10.7554/eLife.33569

Zhou Z, Liu X, Hu Q, Zhang N, Sun G, Cha J, Wang Y, Liu Y, He Q. 2013. Suppression of WC- independent *frequency* transcription by RCO-1 is essential for *Neurospora* circadian clock. Proceedings of the National Academy of Sciences of the United States of America,110(50), E4867–74. doi:10.1073/pnas.1315133110

Zhu Q, Ramakrishnan M, Park J, Belden WJ. 2019. Histone H3 lysine 4 methyltransferase is required for facultative heterochromatin at specific loci. BMC Genomics, 20(1), 1–19. doi:10.1186/s12864-019-5729-7

Zhu Y, Wang X, Forouzmand E, Jeong J, Qiao F, Sowd GA, Engelman AN, Xie X, Hertel KJ, Shi Y. 2018. Molecular Mechanisms for CFIm-Mediated Regulation of mRNA Alternative Polyadenylation. Molecular Cell, 69(1), 62–74. doi:10.1016/j.molcel.2017.11.031

Zimmerman WF, Pittendrigh CS, Pavlidis T. 1968. Temperature compensation of the circadian oscillation in *Drosophila pseudoobscura* and its entrainment by temperature cycles. Journal of Insect Physiology, 14(5), 669–684. doi:10.1016/0022-1910(68)90226-6

